# Cobalamin riboswitches are broadly sensitive to corrinoid cofactors to enable an efficient gene regulatory strategy

**DOI:** 10.1101/2022.02.20.481237

**Authors:** Kristopher J. Kennedy, Florian J. Widner, Olga M. Sokolovskaya, Lina V. Innocent, Rebecca R. Procknow, Kenny C. Mok, Michiko E. Taga

## Abstract

In bacteria, many essential metabolic processes are controlled by riboswitches, gene regulatory RNAs that directly bind and detect metabolites. Highly specific effector binding enables riboswitches to respond to a single biologically relevant metabolite. Cobalamin riboswitches are a potential exception because over a dozen chemically similar but functionally distinct cobalamin variants (corrinoid cofactors) exist in nature. Here, we measured cobalamin riboswitch activity *in vivo* using a *Bacillus subtilis* fluorescent reporter system and found that among 38 tested riboswitches, a subset responded to corrinoids promiscuously, while others were semi-selective. Analyses of chimeric riboswitches and structural models indicate that, unlike other riboswitch classes, cobalamin riboswitches indirectly differentiate among corrinoids by sensing differences in their structural conformation. This regulatory strategy aligns riboswitch-corrinoid specificity with cellular corrinoid requirements in a *B. subtilis* model. Thus, bacteria can employ broadly sensitive riboswitches to cope with the chemical diversity of essential metabolites.

## Introduction

Controlling gene expression is an essential task that cells accomplish in a variety of ways. Non-coding RNAs are one such means of gene regulation, acting in parallel or in concert with historically better-studied protein-based mechanisms (1). In bacteria and archaea, riboswitches are a widespread type of gene regulatory RNA with the distinct ability to sense particular intracellular metabolites by direct binding (2). These RNAs are typically located in the 5’-untranslated region of mRNA transcripts and function as cis-regulators of downstream genes within their transcripts. A riboswitch is composed of an effector-binding aptamer domain and an expression platform. The aptamer domain adopts a three-dimensional structure that can bind its cognate effector molecule. The expression platform domain is a regulatory switch that interprets the effector-binding state of the upstream aptamer typically to promote or disrupt the transcription or translation of downstream genes (3). The diversity of riboswitch effectors and regulatory mechanisms has revealed fundamental insights into how bacteria sense and respond to dynamic environments and has also driven new approaches for precise control and manipulation of microbes for human purposes (4–7).

Cobalamin (Cbl) riboswitches (also called ‘B_12_ riboswitches’ or ‘adenosylcobalamin riboswitches’) are among the most widespread and structurally diverse types of riboswitch in bacteria (8). They directly bind various forms of the enzyme cofactor Cbl as a cognate effector (Figure 1A, C) (9) to regulate genes involved in the biosynthesis, transport, and usage of Cbl. Cbl-dependent enzymes function in common metabolic pathways including methionine synthesis, deoxyribonucleotide synthesis, tRNA modification, and the degradation of certain amino acids, fatty acids, and biopolymers (10–23). Cbl is also required for rarer metabolic processes involved in antibiotic synthesis, mercury methylation, catabolism of steroids, and many others (24–37). Comparative genomic studies indicate that most bacteria perform Cbl-dependent metabolism and that Cbl-riboswitches often regulate these processes (8, 38, 39). However, an overlooked facet among most riboswitch studies is that Cbl is just one member of a class of enzyme cofactors known as corrinoids (Figure 1B) (40). In fact, Cbl-dependent enzymes in bacteria often function with corrinoids other than Cbl. Yet, it remains unclear whether the dozens of naturally occurring corrinoid cofactors are also Cbl-riboswitch effectors (41–47).

**Figure 1.**
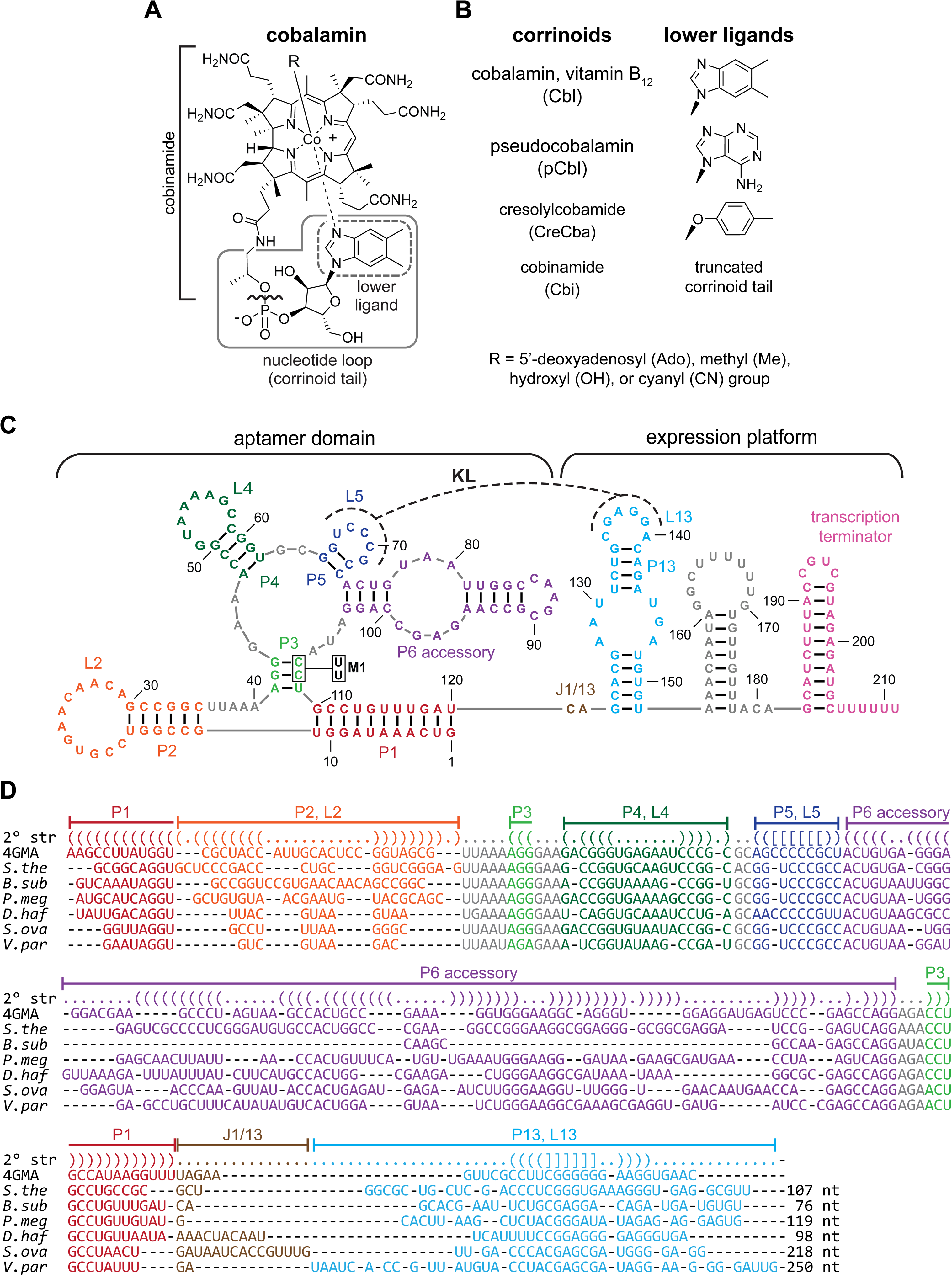
Main corrinoids and riboswitches examined in this study. (A) Chemical structure of cobalamin, also called vitamin B_12_. The gray solid line delineates the corrinoid tail region, which contains the variable lower ligand group in the gray dashed-line box. The R group is the upper ligand. (B) Names, lower ligand structures and abbreviations of the corrinoids used throughout this study. Cbl, pCbl and CreCba are ‘complete corrinoids’ (cobamides). Cbi is an ‘incomplete corrinoid’ that lacks the phosphoribosyl and lower ligand groups as indicated by the bracket and wavy line in the corrinoid tail in panel A. (C) Secondary structural model of the Cbl-riboswitch upstream of *B. subtilis btuF*. Base paired stems (P), loops (L), and junctions (J) are labeled. The P6 accessory region is highly variable in length and number of stems across Cbl-riboswitch sequences, often containing up to six paired regions. In this example, P6 accessory consists of only two paired regions. A kissing loop interaction (KL, dashed line) occurs between L5 of the aptamer and L13 of the expression platform. M1 indicates mutations C107U and C108U, which are examined in Figure 2F. (D) Secondary structural alignments of seven representative Cbl-riboswitches examined in this study. Row “2° str” indicates paired bases as parentheses, loops and junctions as periods, and kissing loop base pairs by brackets. PDB numbers or abbreviations are given for riboswitches from the following organisms and genes: 4GMA, *Thermanaerobacter tencongensis*; *S.the, Symbiobacterium thermophilum cblS*; *B.sub*, *B. subtilis btuF; P.meg*, *Priestia megaterium metE*; *D.haf*, *Desulfitobacterium hafniense* DSY0087; *S.ova*, *Sporomusa ovata btuB2*; *V.par*, *Veillonella parvula mutA*. Non-conserved sequences between P13 and the start codon are indicated by nucleotide sequence length (nt), but the actual sequences were omitted for clarity.

Corrinoid cofactors contain a highly substituted corrin ring with a central cobalt ion, a variable ‘upper ligand’ moiety coordinating the β axial face of the cobalt, and a tail structure extending from the corrin ring and terminating in a variable ‘lower ligand’ moiety that often coordinates the α axial face of the cobalt (Figure 1A, B). The molecular basis of selectivity of Cbl-riboswitches for upper ligand variants of Cbl has been relatively well studied, but selectivity for corrinoid tail variants remains mostly unexplored (9, 48, 49). To our knowledge, only one study has directly examined corrinoid tail specificity of a single Cbl-riboswitch. *In vitro* binding measurements showed that the aptamer of the *Escherichia coli btuB* Cbl-riboswitch binds the complete corrinoids Cbl and 2-methyladeninylcobamide ([2-MeAde]Cba) with a 3.2-fold difference in affinity (*K*_D_ = 89 and 290 nM, respectively). Furthermore, Cbi, an incomplete corrinoid with a truncated tail, binds the aptamer with roughly 8,000-fold lower affinity than Cbl (*K*_D_ = 753 μM), suggesting that this aptamer binds corrinoids in a selective manner (50). In light of these previous studies of corrinoid-specific metabolisms and Cbl-riboswitches, we hypothesize that Cbl-riboswitches harbor a range of distinct corrinoid tail-specific activities.

Here, we examined how a panel of 38 Cbl-riboswitches derived from 12 bacterial species responds to the distinct tail structures of four corrinoids: Cbl, pCbl, and CreCba, representatives of the benzimidazolyl, purinyl, and phenolyl cobamides, respectively, and Cbi, an incomplete corrinoid (Figure 1B). To compare activities among several dozen Cbl-riboswitches, we devised a live cell fluorescence-based reporter system in *Bacillus subtilis*. In contrast to conventional *in vitro* biochemical approaches, this riboswitch reporter system captures the complete corrinoid-responsive gene regulatory process and provides rapid functional measurements with multiple effectors in parallel. Our results obtained from experiments in the reporter system in conjunction with comparative structural analyses of Cbl-riboswitches and corrinoid effectors allowed us to develop a mechanistic model for how corrinoid tail-specific gene regulation is achieved. Additionally, we examine a gene regulatory strategy for the corrinoid specificity of a Cbl-riboswitch and discuss the conceptual and practical implications of these findings.

## Results

### Experimental strategy – Development of an in vivo reporter system

In order to compare corrinoid selectivity among several Cbl-riboswitches and corrinoids, we constructed an *in vivo* GFP reporter system. We initially attempted to use an *E. coli* host for the reporter system but found that most of the riboswitches we tested did not function in the *E. coli* host. We chose *B. subtilis* as an alternative host organism because of the robust genome engineering and gene expression toolsets available. Furthermore, the *B. subtilis* genome does not contain any annotated corrinoid biosynthesis or remodeling genes that would potentially interfere with a riboswitch reporter assay. We engineered the strain to overexpress the Cbl uptake and adenosylation operon, *btuFCDR*, and deleted *queG*, which encodes the only Cbl-dependent enzyme in the genome. We found that *btuFCDR* overexpression increased uptake of not only Cbl, but also pCbl, CreCba and Cbi (Figure 2A-D). Additionally, each corrinoid was transformed from the cyanated to adenosylated form, suggesting that the corrinoids are internalized to the cytoplasm, and not simply accumulating on the outer cell surface. This strain is effective for measuring the response of the *B. subtilis btuF* Cbl-riboswitch to a broad range of concentrations of Cbl (Figure 2E-G). Notably, deletion of *btuR*, which encodes the adensosyltransferase that installs the Ado upper ligand group, rendered the *B. subtilis btuF* Cbl-riboswitch reporter insensitive to exogenously supplied CNCbl, MeCbl, and OHCbl, while retaining dose-dependent repression in response to AdoCbl (Figure S1). This strongly suggests that the *B. subtilis btuF* Cbl-riboswitch only responds to Cbl containing the 5’-deoxyadenosine upper ligand, in contrast with a report suggesting that this riboswitch aptamer can also bind MeCbl and OHCbl (51).

**Figure 2.**
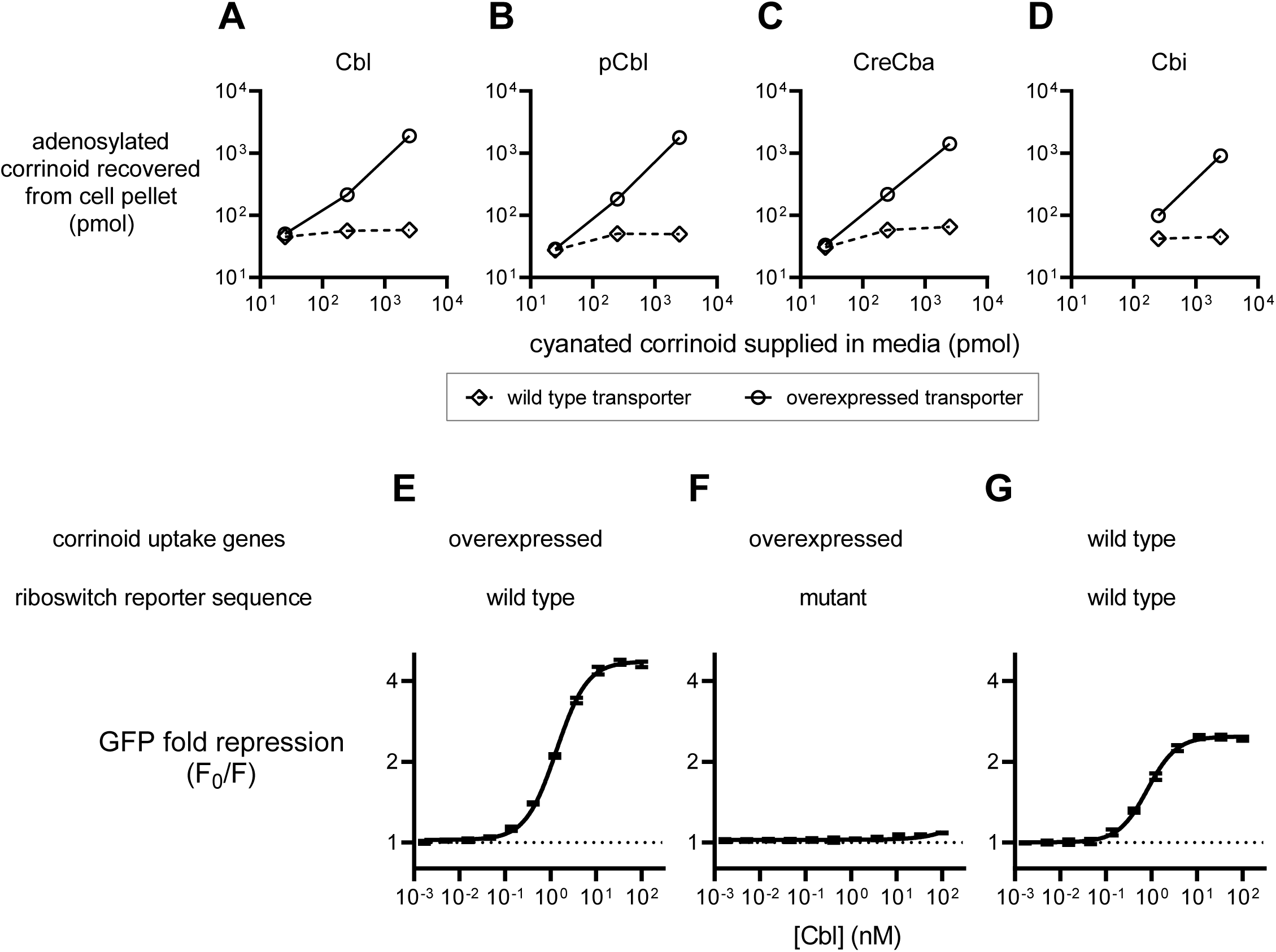
Characterization of a live cell Cbl-riboswitch reporter system. (A-D) Intracellular accumulation of corrinoids in *Bacillus subtilis* strains containing the wild type (diamonds) or constitutively overexpressed (circles) corrinoid uptake genes *btuFCDR*. (E-G) Dose-responses of *B. subtilis btuF* GFP riboswitch reporters. ‘Mutant’ riboswitch refers to the M1 mutant version of the *B. subtilis btuF* riboswitch (Fig. 1C). Data points in panels A-D represent single measurements from one representative experiment. Data points and error bars in panels E-G represent mean and standard deviation of four independent replicates, and horizontal dotted lines demarcate no change in expression.

### Comparison of corrinoid specificity among Cbl-riboswitches

In the strain background described above, we constructed 86 reporter strains to examine riboswitches from 20 bacterial species including 10 species known to produce or require specific corrinoids. Of the 86 reporters, 38 repressed GFP expression 0.5-fold or greater in response to one or more corrinoids. 37 of these 38 functional riboswitch expression platforms contain a predicted intrinsic transcriptional terminator suggesting that they are transcriptional riboswitches. We observed extensive variation in sequence length and nucleotide composition throughout the aptamers and expression platforms of the 38 riboswitches that were functional in *B. subtilis* (Figure 1D). Nine of the functional riboswitches are from *Priestia* (formerly *Bacillus*) *megaterium*, which produces Cbl (41), and 12 are from *Sporomusa ovata* and *Veillonella parvula* which both produce CreCba (44, 45, 52, 53). These results show that Cbl-riboswitches of diverse sequence composition and origin can be examined with the *in vivo* reporter system. To address whether Cbl-riboswitches are corrinoid selective, and how corrinoid selectivity varies among Cbl-riboswitches, we measured the dose-responses of the 38 functional Cbl-riboswitches to four corrinoids, Cbl, pCbl, CreCba and Cbi. Our results show that all of the riboswitches responded to more than one corrinoid, and a subset responded to all four (Figure 3). Strikingly, all of the tested riboswitches are either semi-selective (responding to more than one corrinoid) or promiscuous (responding to all four corrinoids) (Figure 3A-B). We did not find any highly selective riboswitches that respond to only one corrinoid. The semi-selective and promiscuous riboswitches all respond to Cbl and pCbl (Figure 3C), and the promiscuous riboswitches additionally respond to Cbi and CreCba (Figure 3D-E, points above the horizontal dashed line). Furthermore, the semi-selective riboswitches are generally more sensitive to Cbl than to pCbl, while the promiscuous riboswitches respond similarly to these two corrinoids. Almost all of the riboswitches respond weakly to CreCba compared to the other three corrinoids (Figures S2A-C). In general, corrinoid selectivity of a riboswitch appears to be associated with its taxonomic origin (Figure 3C-E). The riboswitches from the Bacilli class are exclusively semi-selective (Figure S2A), whereas those from Negativicutes are predominantly promiscuous (Figure S2B). The *S. ovata cobT* riboswitch is a notable exception that will be discussed later. In contrast to taxonomy, corrinoid selectivity of a riboswitch is not strongly associated with the function of its regulatory target genes (Figure S3).

**Figure 3.**
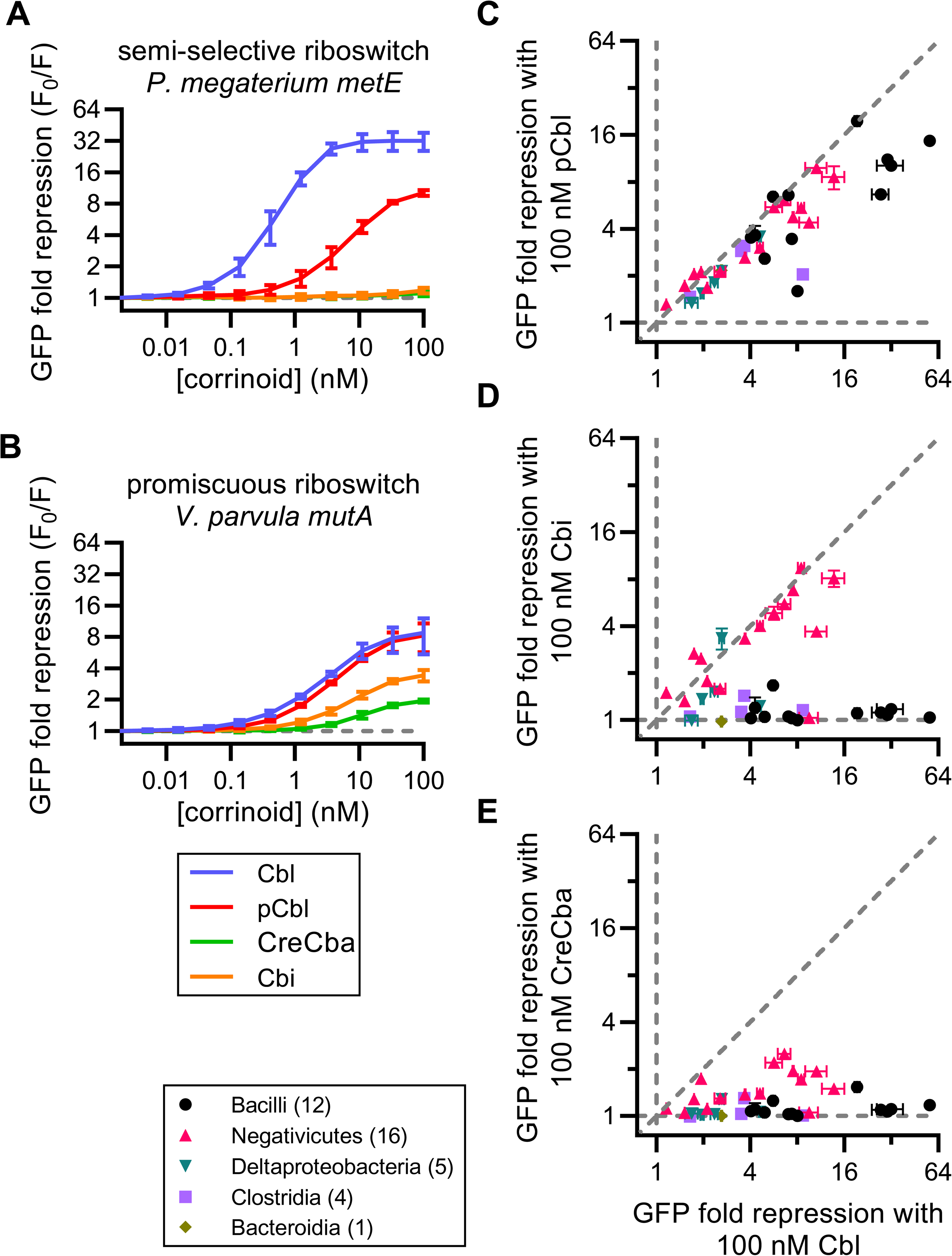
Corrinoid specificity among 38 Cbl-riboswitches. (A,B) Corrinoid dose-responses of two riboswitch reporter strains with distinct corrinoid specificities, representing the semi-selective (A) and promiscuous (B) types. (C-E) Pairwise comparisons of GFP fold repression induced by 100 nM Cbl versus (C) pCbl, (D) Cbi, and (E) CreCba. Data points are colored by taxonomic class with the number of riboswitches analyzed from each class indicated in parentheses. Vertical and horizontal gray dashed lines demarcate a response to only one of the two corrinoids. Diagonal line demarcates equal response to two corrinoids. Data points and error bars in panels A and B represent mean and standard deviation of ten experiments for *P. megaterium metE* and six experiments for *V. parvula mutA*. Data points and error bars in panels C-E represent mean and range of at least two independent experiments for each riboswitch.

Next, we attempted to identify the RNA sequence features that underlie corrinoid selectivity. Chimeric fusions of the semi-selective *P. megaterium metE* riboswitch and the promiscuous *V. parvula mutA* riboswitch enabled us to examine the effects of specific domain and subdomain sequences on corrinoid selectivity (Figure S4). Fusing the *P. megaterium metE* aptamer domain to the expression platform of the *V. parvula mutA* riboswitch produced a semi-selective riboswitch chimera, while the reciprocal chimera was promiscuous, suggesting that the aptamer domain is a major determinant of corrinoid selectivity (Figure S4A). However, results from aptamer subdomain swaps of stem P1, stem-loop P2-L2, stem-loop P4-L4, and P6 accessory region were less conclusive. In the context of the *P. megaterium metE* riboswitch scaffold, swapping stem-loop P2-L2 or the P6 accessory region with the corresponding structures of the *V. parvula mutA* riboswitch produced chimeras that are less selective by gaining sensitivity to Cbi and CreCba (Figure S4B). This suggests that these subdomains confer corrinoid promiscuity. Yet, within the *V. parvula mutA* riboswitch scaffold, swapping stem P1 increased corrinoid selectivity by retaining sensitivity to Cbl and pCbl, but losing sensitivity to CreCba and Cbi (Figure S4C). The remaining chimeras partially or completely lost overall activity. These results demonstrate that subdomains distributed throughout the aptamer domain may impact corrinoid selectivity; no single conserved substructure completely controls corrinoid selectivity, nor did any single structure fully convert a riboswitch’s corrinoid selectivity. Thus, the source of the corrinoid selectivity phenotype appears to be complex and requires inputs from multiple subdomains of the aptamer.

### Corrinoid tail structure impacts selectivity of Cbl-riboswitches

We next sought to identify how structural differences in the corrinoid tail affect the response of semi-selective Cbl-riboswitches to corrinoids. There are no predicted hydrogen bond interactions between the lower ligand and the RNA in the X-ray crystal structures of Cbl-bound riboswitches, making it difficult to surmise how a Cbl-riboswitch might distinguish between corrinoids (51, 54, 55). Could the overall structural conformation of the corrinoid, rather than specific interactions between the RNA and the corrinoid tail, influence Cbl-riboswitch activity?

Corrinoids undergo major conformational changes when spontaneously switching between two distinct states known as ‘base-on’ and ‘base-off’ (56, 57). In the base-on state, a nitrogen atom in the lower ligand base is coordinated to the central cobalt atom of the corrin ring (as shown in Figure 1A). In the base-off state, the lower ligand base is de-coordinated, allowing the tail to move more freely (58, 59). Benzimidazolyl and purinyl cobamides (e.g., Cbl, pCbl) can switch between base-on and base-off states . However, the tail moieties of phenolyl cobamides (e.g., CreCba) and Cbi cannot coordinate cobalt and so these corrinoids exist exclusively in a de-coordinated state (52). Interestingly, we noticed that the semi-selective riboswitches respond strongly to Cbl and pCbl, but respond weakly to Cbi and CreCba. We also observe that semi-selective Cbl-riboswitches are most sensitive to Cbl, which forms the base-on state more readily than pCbl (Figures 3C, S2) (58, 59). In line with these results, the *E. coli btuB* riboswitch aptamer was previously shown to bind Cbl with higher affinity than Cbi and [2-MeAde]Cba (a primarily base-off corrinoid) (50). Also, all six X-ray crystal structures of Cbl-riboswitches contain Cbl in the base-on state (51, 54, 55). Based on these observations, we hypothesized that semi-selective riboswitches distinguish between base-on and base-off states of corrinoids. We therefore used a range of corrinoids with diverse lower ligand structures to test whether the activity of Cbl-riboswitches quantitatively correlates with the base-on tendency of corrinoids.

We selected a panel of sixteen corrinoids for this analysis, including both natural and synthetic benzimidazolyl, purinyl, and aza-benzimidazolyl corrinoids. These corrinoids span a range of base-on tendency between that of Cbl and pCbl, which we measured as the ratio of spectral absorbance at 525 and 458 nm in the adenosylated form (Figures 4A, S5). Base-on/base-off equilibrium constants for AdoCbl, Ado[2-MeAde]Cba, and pCbl in aqueous conditions have been reported as 76, 0.48, and 0.30, respectively, and are consistent with our measurements of corrinoid base-on tendency (58, 59). We observed a strong association between base-on tendency and riboswitch response in semi-selective riboswitches (Figure 4B). Even among promiscuous Cbl-riboswitches, we observe measurable sensitivity to base-on tendency, albeit to a much smaller degree (Figure 4C). These results support the hypothesis that Cbl-riboswitches selectively respond to corrinoids by distinguishing between the base-on and base-off states of corrinoids.

**Figure 4.**
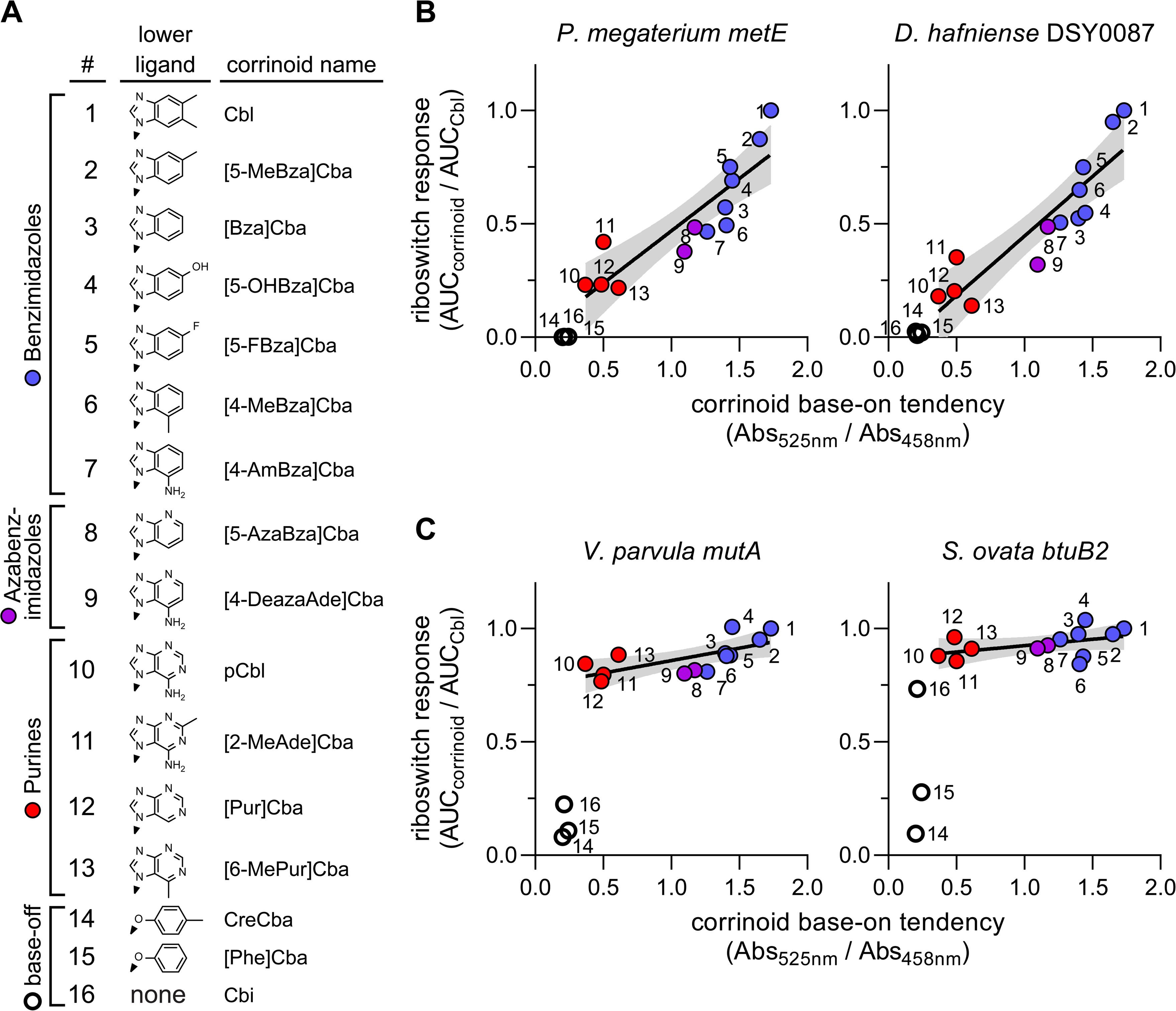
Corrinoid base-on tendency correlates with corrinoid potency. (A) Sixteen adenosylated corrinoids used to test the relationship between corrinoid base-on tendency and corrinoid potency. (B) Semi-selective riboswitches *P. megaterium metE* and *D. hafniense* DSY0087 and (C) promiscuous riboswitches *V. parvula mutA* and *S. ovata btuB2* were used to measure the response to each corrinoid. Base-on tendency was measured as the ratio between spectral absorbance at 525 and 458 nm in a pH 7.3 solution. Absorbance spectra are displayed in Figure S5. The cumulative corrinoid response was measured as the area under the dose-response curve of the riboswitch reporter strain for that corrinoid (AUC_corrinoid_) relative to its dose-response to Cbl (AUC_Cbl_). Corrinoids with benzimidazole (blue circles), azabenzimidazole (purple circles), and purine (red circles) lower ligands can adopt the base-on conformation. The corrinoids that are unable to adopt the base-on state (CreCba, [Phe]Cba and Cbi) are represented by empty circles. Trendlines in B and C were fit to data points of corrinoids with benzimidazole, azabenzimidazole and purine lower ligands, with strictly base-off corrinoids excluded. Each data point represents a single measurement of riboswitch dose-response and corrinoid base-on tendency.

### Structural comparisons between base-on and base-off tail orientations

The results presented above led us to speculate about how a Cbl-riboswitch could detect the base-on and base-off state of a corrinoid. In all published X-ray crystal structures of Cbl-riboswitches, aptamer-effector binding is achieved mainly through van der Waals forces and shape complementarity between the binding site and base-on Cbl. Only a few hydrogen bonds between the RNA and corrinoid are observed, none of which occur with the lower ligand group of Cbl (51, 54, 55). Thus, it appears unlikely that the riboswitch is directly detecting the specific chemical differences among corrinoid lower ligands. Instead, we considered whether the riboswitch discriminates base-on and base-off forms of a corrinoid by sensing corrinoid conformation. In the base-on state, the tail is spatially constrained due to the Co-N coordinate bond, whereas in the base-off form it is able to sample a wider range of spatial positions (60). To develop mechanistic insight into how the base-on and base-off states a corrinoid could impact Cbl-riboswitch activity, we leveraged the plethora of publicly accessible X-ray crystal structures of macromolecule-bound Cbl (37, 51, 54, 55, 61–70). We first assessed the range of structural conformations that are potentially sampled by corrinoids as they dynamically switch between base-on and base-off states by aligning and visually comparing various structural models of Cbl. Six base-on Cbl models were obtained from structural studies of Cbl-riboswitches and synthetic Cbl RNA aptamers, whereas base-off/His-on Cbl models were obtained from X-ray crystal structures of ten Cbl-dependent enzymes (Table S2). After aligning and superimposing these molecular models by their central cobalt and coordinating nitrogen atoms, we observed that the corrin rings and their amide and methyl substituents occupy similar spatial positions, but the tails of base-on and base-off Cbl structures occupy distinct positions (Figure 5A-B). Moreover, the base-on Cbl tails appear in very similar positions with lower ligands in close proximity to the central cobalt ion, whereas the base-off Cbl tails appear more scattered with lower ligands more distal to the cobalt ion. These structural alignments visually convey the degree to which the conformations of base-on and base-off corrinoids can vary among biomolecular complexes.

**Figure 5.**
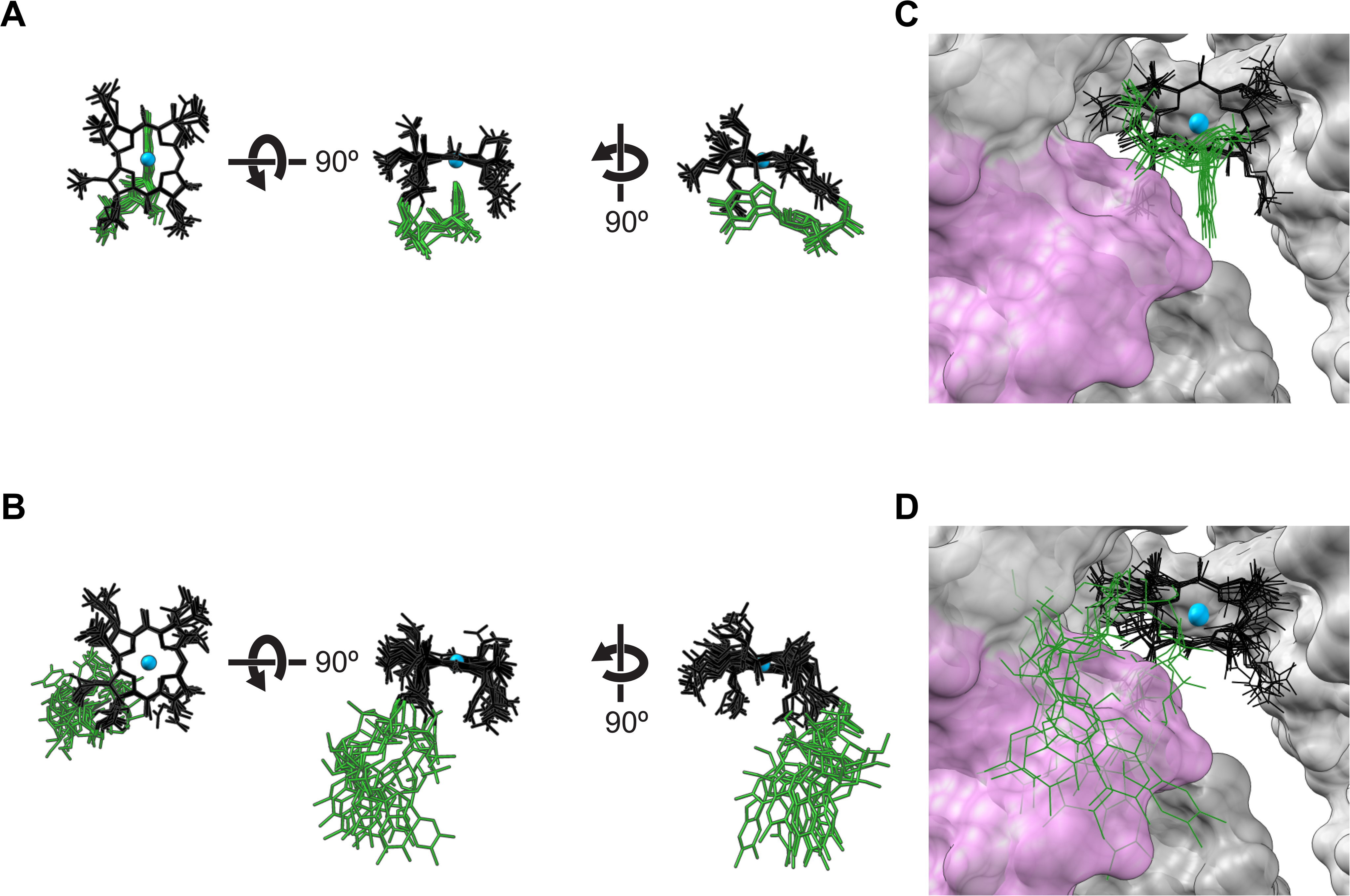
Distinct tail positions among base-on and base-off corrinoids may impact binding of corrinoids to riboswitches. 3D alignments of Cbl structural models derived from published X-ray crystal structures of (A) base-on Cbl in complex with RNAs and (B) base-off Cbl in complex with proteins. The Cbl-binding site in the X-ray crystal structure of the *Thermoanaerobacter tengcongensis* Cbl-riboswitch (PDB ID 4GMA) is depicted with (C) base-on and (D) base-off Cbl alignments. Cbl models were aligned by the cobalt and coordinating nitrogen atoms in the corrin ring. Structures of the corrin ring, cobalt, and tail of Cbl are colored in black, blue, and green, respectively. Upper ligand structures of Cbl were omitted for clarity. Riboswitch RNA structures are depicted as space-filled models with the L5-L13 kissing loop in pink and the rest of the RNA in gray.

Next, we compared the positions of the aligned base-on and base-off Cbl models in the context of 3D Cbl-riboswitch models. We analyzed X-ray crystal structures of the two Cbl-riboswitches that contain resolved kissing loop structures: one from *Thermoanaerobacter tengcongensis* (Figure 5C-D) and one identified from a marine metagenome sequence (Figure S6A-B) (54). These structural models show that the base-on tails are contained within the binding site, whereas the tails of the base-off Cbl structures protrude away from the binding site and clash with the L5-L13 kissing loop. Although the *B. subtilis btuF* (Figure S6C-D) and *Symbiobacterium thermophilum cblT* (Figure S6E-F) riboswitch models do not contain the L13 structure of the kissing loop, some of the modeled base-off tails clash with L5 of the aptamer in these structures (51, 55). The kissing loop has been shown to play a key mechanistic role of sensing the corrinoid-binding state of the aptamer domain to influence downstream regulatory structures in the expression platform (71, 72). If kissing loop formation in semi-selective Cbl-riboswitches is sensitive to the base-on and base-off states of the corrinoid, then corrinoid selectivity may be mediated by either selective binding by the aptamer or selective formation of downstream regulatory structures.

To determine whether the expression platform structures can impact corrinoid selectivity, we examined corrinoid-selective binding separately from subsequent corrinoid-selective regulation. We tested for promiscuous binding by comparing the Cbl dose-response of the *P. megaterium metE* Cbl-riboswitch in the presence and absence of competing 100 nM Cbi and found that the response to Cbl is unaffected by Cbi (Figure S7A-B). This indicates that Cbi does not compete with Cbl for riboswitch binding, supporting corrinoid-selective binding as the mechanism of semi-selectivity. However, when the aptamer of this semi-selective riboswitch is replaced with the aptamer of the promiscuous *S. ovata nikA* riboswitch, it retains semi-selectivity, which suggests that the *P. megaterium* expression platform also plays a role in corrinoid selectivity (Figure S7A). Interestingly, the Cbl dose-response of the *S. ovata nikA* / *P. megaterium metE* chimeric riboswitch does become sensitized to competing Cbi addition, confirming that the *S. ovata nikA* aptamer retains sensitivity to base-off corrinoid in the context of this chimera (Figure S7C). Taken together, these results show that base-off corrinoids may impede both Cbl-riboswitch binding and formation of regulatory structures, explaining the link between corrinoid base-on tendency and riboswitch activity observed in Figure 4.

### Gene regulatory strategy of corrinoid selectivity

While the prior experiments clearly demonstrate that Cbl-riboswitches are capable of distinguishing between corrinoids, we wondered what purpose Cbl-riboswitch corrinoid selectivity might serve in the organisms containing these regulatory systems. We posit that Cbl-riboswitch selectivity reflects a regulatory strategy that complements the corrinoid-specific requirements of the cell and avoids gene mis-regulation. As a specific example, we hypothesize that only corrinoids that are functionally compatible with a Cbl-dependent enzyme should cause riboswitch-mediated repression of the expression of its Cbl-independent counterpart (Figure 6A, B). We tested this hypothesis directly in *B. subtilis* by examining the function and regulation of methionine synthase isozymes MetE (Cbl-independent) and MetH (Cbl-dependent). In bacterial genomes with both *metE* and *metH*, a tandem SAM-Cbl riboswitch is commonly found upstream of the *metE* gene (73, 74). The *B. subtilis* genome contains *metE* but lacks *metH*, and no Cbl-riboswitch is located upstream of *metE*. We therefore constructed strains of *B. subtilis* that heterologously express the *metE* or *metH* locus from the Cbl-producing species *P. megaterium*, in a Δ*metE* background with overexpressed corrinoid uptake genes (Figure 6C) (75–77). In each strain, the *P. megaterium* genes are constitutively transcribed from the promoter P_Veg_ and also contain a transcriptionally fused *gfp* to measure expression levels. Growth of the *B. subtilis* strain expressing *P. megaterium metH* in a medium lacking methionine was supported to varying extents by most benzimidazolyl and both phenolyl cobamides, but not by [5-OHBza]Cba, the purinyl cobamides, or Cbi (Figure 6D, red squares). This result indicates that MetH-dependent growth is influenced by the corrinoid tail structure, as observed previously in other bacteria (78–82). In the *B. subtilis* strain containing the *P. megaterium metE* locus which includes the repressing SAM-Cbl-riboswitch, growth was suppressed by benzimidazolyl cobamides to different extents (Figure 6D, blue circles). This growth pattern coincides with the GFP repression measured for the *metE* riboswitch (Figure 6E), except that the repression of *metE* by purinyl cobamides is apparently insufficient to suppress growth in this context. Comparison of the two strains in response to a suite of corrinoids reveals a striking correspondence between riboswitch-mediated suppression of growth in the *metE*-containing strain and growth promotion by corrinoids in the *metH*-containing strain (Figure 6C). [5-OHBza]Cba and the phenolyl cobamides are exceptions to the trend, though in neither case is MetE-dependent growth completely suppressed by a corrinoid incompatible with MetH. This result demonstrates that riboswitch-based repression and cobalamin-dependent isozyme function are largely aligned for the *P. megaterium metE-metH* pair and suggests that Cbl-riboswitch specificity may generally adhere to a regulatory strategy reflecting the cell’s corrinoid preference.

**Figure 6.**
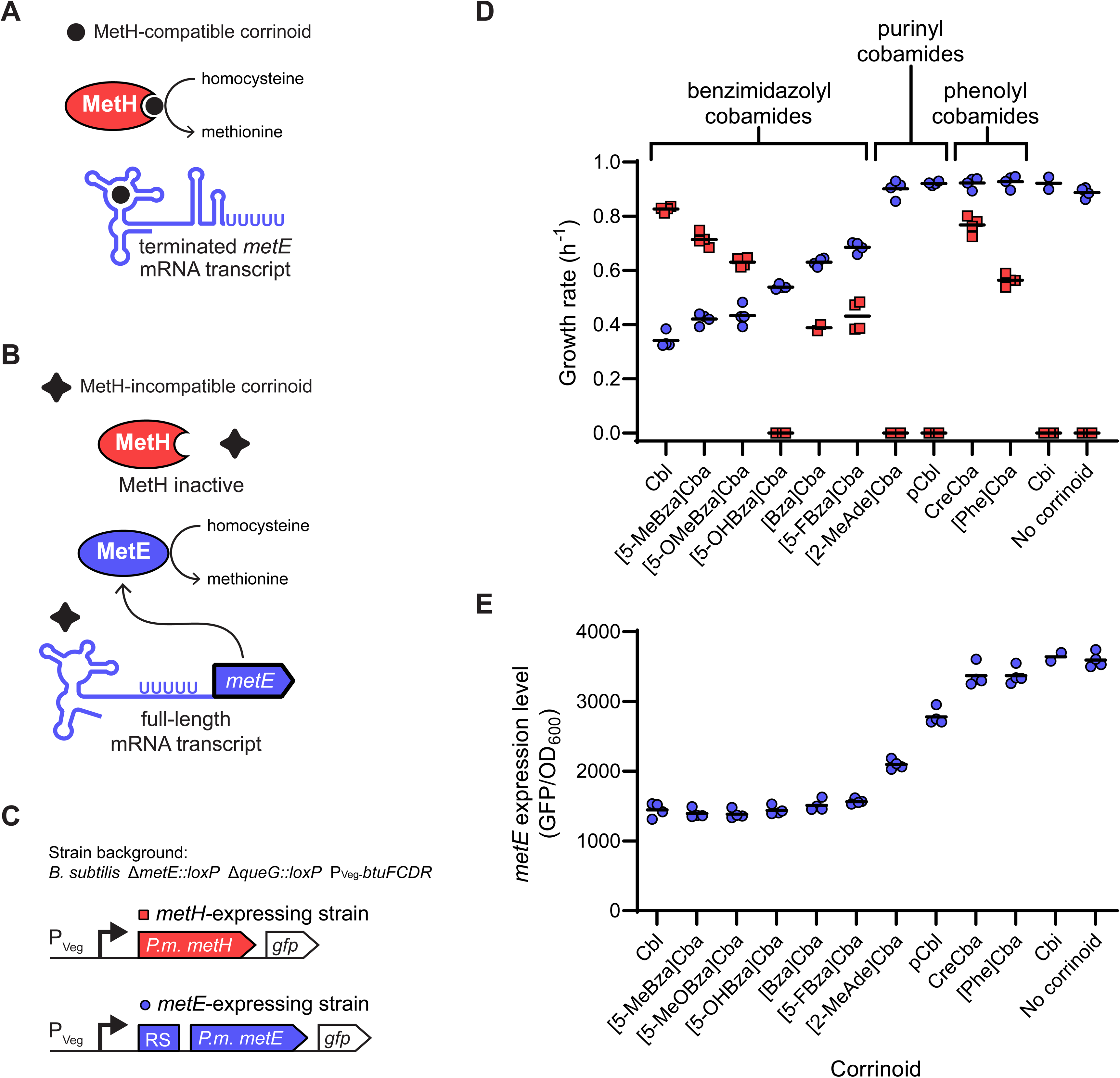
Corrinoid specificities of the *P. megaterium metE* riboswitch and *P. megaterium* MetH enzyme are aligned. Expression of the Cbl-independent methionine synthase MetE and enzymatic activity of the Cbl-dependent methionine synthase MetH are potentially impacted by the presence of (A) MetH-compatible corrinoids and (B) MetH-incompatible corrinoids. (C) The *metH*-expressing strain (red) is a *B. subtilis* Δ*metE::loxP* Δ*queG::loxP* P_Veg_-*btuFCDR* strain heterologously expressing *P. megaterium metH*. The *metE*-expressing strain (blue) is a *B. subtilis* Δ*metE::loxP* Δ*queG::loxP* P_Veg_-*btuFCDR* strain heterologously expressing *P. megaterium metE* downstream from a SAM-Cbl tandem riboswitch (RS). Each methionine synthase gene is constitutively expressed from the P_Veg_ promoter and is transcriptionally fused to *gfp*. (D) Growth rates of the *metE*-expressing strain (blue circles) and *metH*-expressing strain (red squares) strains were measured in methionine-dependent culture conditions containing 20 nM corrinoid. (E) Expression of *metE* was measured as GFP fluorescence per OD_600_ in medium lacking methionine, supplemented with 20 nM corrinoids. Data points are individual measurements and black horizontal lines represent the mean of the four replicate measurements.

## Discussion

Riboswitches are key regulators of microbial gene expression. The Cbl-riboswitch was the first type discovered and is among the most widely distributed riboswitch classes in bacteria and archaea (9, 83). Previous biochemical and structural studies have uncovered the major molecular features of the Cbl-riboswitch response to Cbl, including how upper ligand variants of Cbl impact their function (48, 49, 51, 54, 55, 71, 72, 74, 84). Yet few studies have examined how other naturally occurring corrinoids containing diverse lower ligand structures impact gene regulation by Cbl-riboswitches (50). Here, we found that Cbl-riboswitches vary in their ability to discriminate between corrinoids, with some being semi-selective on the basis of corrinoid base-on/off state, and others being promiscuous. These results were enabled by a carefully designed fluorescent reporter system capable of measuring the responses of dozens of Cbl-riboswitches to multiple corrinoids *in vivo* (Figures 2, S1). Since several naturally occurring corrinoids other than Cbl appear to be potent effectors for Cbl-riboswitches, we propose that the term ‘corrinoid riboswitch’ be adopted to describe this broad class of RNAs more accurately. This may also mitigate the inconsistent and overlapping terminology used in the literature (i.e. cobalamin riboswitch, adenosylcobalamin riboswitch, B_12_ riboswitch, vitamin B_12_ riboswitch, etc.).

We can roughly estimate the intracellular corrinoid concentrations in our corrinoid dose-response experiments to assess the physiological relevance the *in vivo* Cbl-riboswitch data. With the observation that most of the corrinoid in the medium is imported by the *B. subtilis* riboswitch reporter cells (Figure 2A-D), and assuming a cell volume of 10^-15^ L and a culture cell titer of 10^12^ cells/L at OD_600_ = 1, we calculate the cytoplasmic corrinoid concentrations to range from 0.01 to 100 μM in the corrinoid dose-response experiments – a 1,000-fold increase from the cell culture medium to the cytoplasm. Reports of binding affinity (*K*_D_) for riboswitch aptamers to AdoCbl range from 0.026 to 90 μM, which is within the range of our estimated intracellular corrinoid concentrations (48, 50, 54, 72, 85). Additionally, Cbl uptake has been measured in some bacterial species. In *E. coli*, the minimum cytoplasmic Cbl concentration to support MetH-dependent growth is roughly 0.03 μM (86). In studies of Cbl uptake across several bacterial species, saturating Cbl uptake can result in cytoplasmic Cbl concentrations in the low μM to low mM range, depending on the species (87, 88). Taken together, these data and calculations support the physiological relevance of the Cbl-riboswitch responses measured in this study.

We observed that Cbl-riboswitches display different degrees of corrinoid selectivity, with some that respond to a subset of corrinoids (semi-selective) and others that respond to all tested corrinoids (promiscuous) (Figures 3, S2). Our chimeric riboswitch results suggest that sequence and structural determinants of corrinoid selectivity are dispersed throughout the Cbl-riboswitch aptamer scaffold rather than being confined to a single conserved region (Figure S4). This finding contrasts with other studies of riboswitch specificity. For example, in a study of Cbl upper ligand specificity, a few key residues in the Cbl binding site were sufficient to fully convert a Cbl-riboswitch from MeCbl-specific to AdoCbl-specific (48). In purine riboswitches, effector specificity is achieved by positioning of a critical conserved uracil or cytosine residue in the binding site of the aptamer, which forms a base-pair with the adenine or guanine effector, respectively. Among the various families of SAM riboswitches, highly specific binding of SAM and exclusion of SAH is achieved by RNA structures that discriminate the charged sulfonium ion of SAM from the uncharged sulfoether of SAH (89). In contrast, the SAM/SAH riboswitch class attains effector promiscuity for SAM and SAH by a general lack of interaction between the RNA and the aminocarboxypropyl side chains of these effectors (90, 91). This is reminiscent of Cbl-riboswitches which similarly have few molecular contacts between the RNA and corrinoid tail (50, 54, 55, 84).

Overall, our structural model analyses support a mechanism in which semi-selective Cbl-riboswitches primarily sense the distinct corrinoid tail orientations of the base-on and base-off forms. In this case, Cbl-riboswitches only indirectly sense the chemical composition of the variable lower ligand group, with differential binding largely determined by steric effects and shape complementarity (Figures 4-5). Recent molecular dynamics simulations of the *T. tengcongensis* Cbl-riboswitch suggest that the kissing loop structure may form prior to effector binding, which would place even greater constraints on the corrinoid tail orientation to achieve shape complementarity with its binding site (92). Additionally, some Cbl-riboswitches may in fact bind base-off corrinoids but disrupt subsequent formation of downstream regulatory structures of the expression platform, perhaps by interfering with the kissing loop (Figure S7). A similar feature has been observed in tetrahydrofolate (THF) riboswitches where chemical variations in the para-aminobenzoic acid moiety of THF analogs differentially perturb expression platform structures without affecting aptamer binding (93). The mechanisms of corrinoid selectivity of Cbl-riboswitches could be directly tested in future structural or biochemical studies of promiscuous Cbl-riboswitches with base-on and base-off corrinoids.

In regard to gene regulatory strategies, it seems sensible that Cbl-riboswitches are not highly effector-specific because bacteria are often flexible in their corrinoid usage. A variety of corrinoids have been shown to support growth of *C. difficile*, *S. ovata, and Ensifer meliloti* despite each of these organisms displaying highly specific corrinoid production (82, 94, 95). Furthermore, since corrinoid auxotrophy is prevalent among corrinoid-dependent bacteria, many organisms may need to take advantage of the wide range of corrinoids that may be available in their environment (96). Thus, the range of effector selectivity that we observe among Cbl-riboswitches may reflect a coevolution between corrinoid-responsive gene regulation and corrinoid-dependent physiology. Our result demonstrating complementary corrinoid selectivity between *P. megaterium* MetH-dependent growth and Cbl-riboswitch-dependent expression of MetE is consistent with this notion (Figure 6).

Alternatively, the preference for base-on corrinoids among Cbl-riboswitches may function as a proxy to discriminate complete corrinoid coenzymes from incomplete corrinoids such as Cbi, which often function poorly as coenzymes. This idea has been proposed as an explanation for the remarkably high selectivity of the corrinoid uptake system in mammals (97). Interestingly, we found that all but one of the *S. ovata* riboswitches tested are promiscuous types that can respond to its natively produced CreCba. The notable exception is a semi-selective riboswitch upstream of the gene *cobT* (Figure S2B), which functions in a late step of corrinoid biosynthesis that occurs after synthesis of Cbi (53, 98–103). Thus, this Cbl-riboswitch that discriminates against Cbi may allow homeostatic regulation of *cobT* in response to complete corrinoids like Cbl, while preventing unproductive repression of *cobT* in the presence of incomplete corrinoids like Cbi. Future studies examining corrinoid-specific gene regulation of riboswitches in the context of their native organisms may help clarify which regulatory strategies are generally at play in corrinoid-related bacterial physiology.

Our findings fit into a broader discussion of how corrinoids impact complex microbial communities (104). In future studies, it will be worth examining how the interplay between corrinoid-specific gene regulation and corrinoid-specific metabolic pathways influence microbial interactions. There is also significant interest in the fields of bioengineering and synthetic biology to use riboswitches as gene regulatory devices because they act more rapidly and efficiently than protein-based regulatory systems (6, 7, 105, 106). Riboswitches have also become desirable therapeutic drug targets because they often control essential metabolic pathways in pathogenic microbes (4, 5, 107). Broader consideration of the conformational dynamics of larger types of effector molecules including organic cofactors, antibiotics, and their analogs could aid efforts to engineer synthetic RNA-based regulatory platforms that function robustly *in vivo*. This could also inform future efforts to develop synthetic antimetabolites that, for example, elicit gene mis-regulation, instead of a more common strategy of creating inhibitors for enzymes (108, 109).

The chemical diversity of corrinoids is intrinsically linked to a vast array of metabolic processes and microbial interactions. Yet it remains unclear how microbes have evolved to cope with and thrive on the assortment of natural corrinoid analogs, especially when compared to other primary metabolites including organic cofactors, nucleotides and amino acids which typically require one specific structural form for precise biological functions. We have gained new appreciation for the impacts of chemical diversity on biological function by focusing on the Cbl-riboswitch with its distinctively complex structure and regulatory mechanism, and by accounting for the often overlooked biological and ecological roles of corrinoid analogs. Future studies into the evolution of microbial molecular specificity for corrinoids may yield further insight into the nature of these exceptionally versatile coenzymes.

## Materials and methods

### Cbl-riboswitch sequence analysis

Cbl-riboswitch sequences, chromosomal coordinates and regulon information were downloaded from the RiboD online database (73) (Table S1). The genome of *Sporomusa ovata* was not included in the RiboD database, so we used the RiboswitchScanner webserver to search for Cbl-riboswitches in this organism (110, 111). Cbl-riboswitch aptamer sequences were manually aligned by conserved secondary structures bounded by the 5’ and 3’ ends of the P1 stem (8, 85) (Data S1). The P13-L13 stem loop and potential intrinsic transcriptional termination hairpin structures of the expression platform were identified using secondary structure prediction tools in RNAstructure 6.2 (112, 113). Intrinsic terminators were identified as stem loops directly preceding a sequence of five or more consecutive uracil residues (114, 115). Sequence alignment and annotation was carried out in JalView 2.11.1.4 (116). Cartoons of riboswitch secondary structures were constructed using the StructureEditor program of RNAstructure 6.2 (113).

### Corrinoid production, extraction, purification, and analysis

Cyanocobalamin, adenosylcobalamin, methylcobalamin, hydroxocobalamin, and dicyanocobinamide were purchased from MilliporeSigma. All other corrinoids used in this study were produced in bacterial cultures and purified in cyanated form as previously described (82, 95, 117, 118). For the experiments in Figures 4 and S5, corrinoids other than cobalamin were chemically adenosylated to obtain the coenzyme (5’-deoxyadenosylated) form as previously described (82, 117).

UV/Vis spectra were collected from corrinoid samples in UV/Vis-transparent 96-well microtiter plates (greiner bio-one UV-STAR® 675801) using a BioTek Synergy 2 or Tecan Infinite M1000 Pro plate reader. To measure concentrations of corrinoid stock solutions, corrinoid samples were diluted 10-fold in 10 mM sodium cyanide to obtain the dicyanated base-off form of the corrinoid. The concentration of the dicyanated corrinoid was calculated using the extinction coefficient ε580 = 10.1□mM^−1^□cm^−1^ (52, 119). For adenosylated corrinoids used in Figures 4 and S5, base-on tendency at neutral pH was measured as the ratio of spectral absorbance at 525 nm and 458 nm in phosphate buffered saline solution pH 7.3 at 37°C (47, 59).

### Plasmid and strain construction

Plasmids generated in this study were constructed with one-step isothermal assembly (120) and introduced into *E. coli* strain XL1-Blue by heat shock transformation. Riboswitch reporter plasmids were constructed in the shuttle vector pSG29 for single copy integration at the amyE locus of the *B. subtilis* chromosome (121). Riboswitch DNA sequences were inserted between the transcriptional start site of the constitutive P_Veg_ promoter and the *gfp* translational start site of pSG29. For riboswitch sequences that resulted in no detectable GFP signal under any conditions, a synthetic ribosome binding site (RBS) sequence R0 (5’-GATTAACTAATAAGGAGGACAAAC-3’) from pSG29 was placed between the riboswitch sequence and *gfp* translational start site (Table S1B-C).

Strains used in this study are listed in Table S1A. All *B. subtilis* riboswitch fluorescent reporter strains and *B. subtilis* strains expressing *P. megaterium metE* and *metH* used in this study are derived from the high-efficiency transformation strain SCK6, which has a xylose-inducible competence gene cassette (122). Preparation of competent cells and transformations of all SCK6-derived strains were performed as previously described (122). The strain KK642, which constitutively overexpresses the corrinoid uptake genes, was constructed by deletion of gene *queG* and replacement of the promoter and 5’ untranslated region of the *btuFCDR* operon with the P_Veg_ promoter and R0 RBS (121). *B. subtilis* genes *queG*, *btuR*, and *metE* were targeted for deletion by recombination with kanamycin resistance cassettes containing flanking sequence homology to each respective locus. Kanamycin resistance cassettes were PCR-amplified from genomic DNA of *B. subtilis* strains BKK08910 (Δ*queG*::*kan^R^*), BKK33150 (Δ*btuR*::*kan^R^*), and BKK13180 (Δ*metE*::*kan^R^*) (123). Kanamycin resistance cassettes were removed by Cre-Lox recombination using plasmid pDR244 as previously described (123).

*B. subtilis* strains heterologously expressing *metE* and *metH* from *P. megaterium* were constructed as follows. The *metE* and *metH* genes were PCR-amplified from genomic DNA of *P. megaterium* DSM319 and cloned between the transcriptional start site and the *gfp* translational start site of pSG29. The *P. megaterium metE* amplified fragment starts at the SAM-Cbl tandem riboswitch in the 5’ UTR and ends at the *metE* stop codon, whereas the *P. megaterium metH* fragment starts at the *metH* RBS, which is composed of 20 nucleotides preceding the *metH* translational start site and ends at the *metH* stop codon.

Riboswitch reporter plasmids and plasmids containing *P. megaterium metE* and *metH* were linearized by restriction enzyme digest with ScaI-HF (New England Biolabs) and selected for integration at the *amyE* locus of *B. subtilis* by plating on lysogeny broth (LB) agar with 100 µg/mL spectinomycin. Colonies were screened for integration at *amyE* by colony PCR. All stocks of bacterial strains were stored in 15% glycerol at -80 □C.

### Intracellular corrinoid accumulation experiments

*B. subtilis* strains were inoculated from single colonies into 50 mL LB and grown with aeration at 37 □C in a shaking incubator (Gyromax 737R, Amerex Instruments, Inc.) for 5-6 hours until reaching an optical density at 600 nm (OD_600_) of 1.0 to 1.5. Each culture was diluted 10-fold in LB and split into 13 25 mL cultures containing 0, 25, 250, or 2500 picomoles of a cyanated corrinoid (Cbl, pCbl, CreCba and Cbi). These cultures were incubated at 37 □C with aeration for 3-4 hours to a final OD_600_ of 1.5 to 2.0. The cells were pelleted by centrifugation at 4,000 g for 10 min. Cell pellets were rinsed three times by resuspension in 10 mL phosphate buffered saline solution pH 7.3 followed by centrifugation. After the final centrifugation, tubes were wrapped in aluminum foil to protect adenosylated corrinoids from exposure to light.

To extract intracellular corrinoids, cell pellets were resuspended in 5 mL of 100% methanol by vigorous vortexing for 30 secs. Samples were stored at -80 □C until the next day. Frozen lysates were heated in an 80 □C water bath for 1.5 hours, with 15 seconds of vortexing every 30 minutes. Methanol concentration of each sample was diluted to 10% by adding 45 mL of water, and cell debris was pelleted by centrifugation at 4,000 g for 10 mins. The supernatants were used for the subsequent steps.

All of the following steps were carried out in darkened rooms illuminated with red light to preserve light-sensitive adenosylated corrinoid samples. Solid-phase extraction of adenosylated corrinoids with Sep-Pak C18 cartridges (Waters) was performed as previously described (82).

Solvents were evaporated in a vacuum concentrator centrifuge (Savant SPD1010, Thermo Scientific) at 45 □C and the samples were resuspended in 500 µL deionized water and passed through 0.45 µm pore-size filters (Millex-HV, MilliporeSigma).

Corrinoids were analyzed on an Agilent 1200 series (high-performance liquid chromatography system equipped with a diode array detector (Agilent Technologies). Samples were injected into an Agilent Zorbax SB-Aq column (5-μm pore size, 4.6 by 150□mm). The following HPLC method was used: solvent A, 0.1% formic acid–deionized water; solvent B, 0.1% formic acid–methanol; flow rate of 1□mL/minute at 30°C; 25% to 34% solvent B for 11□minutes, followed by a linear gradient of 34% to 50% solvent B over 2□minutes, followed by a linear gradient of 50% to 75% solvent B over 8 minutes.

### Riboswitch fluorescent reporter assays

Corrinoid dose-response assays of riboswitch reporter strains were set up as follows. Saturated cultures of the riboswitch reporter strain in LB were diluted 200-fold in LB and dispensed into 96-well microtiter plates (Corning Costar Assay Plate 3904) containing a range of concentrations of various corrinoids. The plates were sealed with gas diffusible membranes (Breathe-Easy, MilliporeSigma) and incubated at 37 □C for 4 to 5 hours in a benchtop heated plate shaker (Southwest Science) at 1,200 revolution per minute (rpm). GFP fluorescence (excitation/emission/bandwidth = 485/525/10 nm) and absorbance at 600 nm (A_600_) were measured on a Tecan Infinite M1000 Pro plate reader. The A_600_ measurements of uninoculated medium and fluorescence measurements of the parental control strains lacking *gfp* were subtracted from all readings. Data were plotted and analyzed in GraphPad Prism 9.

### 3D structural analysis of corrinoids and macromolecular models

Molecular models of cobalamin in the base-on and base-off/His-on state in complex with various proteins and RNAs were downloaded from the Protein DataBank (PDB) (Table S2) (124). PDB files were analyzed in UCSF Chimera 1.14 (125). Corrinoid molecular models were aligned with each other by the central cobalt atom and coordinating nitrogen atoms of the corrin ring, using the PDB ID 4GMA Cbl model as a reference. Corrinoid models were aligned within the binding sites of riboswitch structures PDB IDs 4GMA, 4FRN, 4GXY, and 6VMY (51, 54, 55).

### Methionine-dependent growth of *B. subtilis* strains

*B. subtilis* strains were streaked from frozen stocks onto LB agar plates and incubated overnight at 37 □C for 14-18 hrs. Single colonies were used to inoculate 3 mL liquid starter cultures containing Spizizen minimal medium supplemented with 0.02% D-glucose and 0.2% L-Histidine (SMM) (126). Starter cultures of the *metH*-expressing strain were supplemented with 1 nM CNCbl to support growth, whereas the *metE*-expressing strains were cultured in SMM without CNCbl. Starter cultures were incubated overnight shaking (250 rpm, 37 □C) for 20 hours, reaching cell density of about OD_600_ = 1.0. Starter cultures were diluted 500-fold by transferring 50 μL of starter culture to 25 mL of SMM. Then 75 μL of the diluted culture were dispensed into wells of a 96 well microtiter plate containing 75 μL of SMM supplemented with 40 nM of various corrinoids. Plates were sealed with gas diffusible membranes (Breathe-Easy, MilliporeSigma) and incubated at 37 □C on the ‘high shaking’ setting of a BioTek Synergy2 plate reader. Growth kinetics and *metE* and *metH* expression were measured by A_600_ and GFP fluorescence every 15 minutes for 72 hours. Data were plotted and analyzed in GraphPad Prism 9.

## Acknowledgments

We thank past and present members of the Taga lab for support. In particular, we thank Sebastian Gude, Zachary Hallberg, and Gordon Pherribo for critical reading of the manuscript; Alexa Nicolas and Amanda Shelton for helpful discussions; Anna Grimaldo and Jong-Duk Park for reagents. We thank Amrita Hazra, Arash Komeili, Michael Marletta, Kathleen Ryan, and Matthew Traxler for helpful discussions. We especially thank Ming Hammond for critical reading of the manuscript and helpful advice. We are grateful to the labs of Benjamin Blackman, John Deuber, Krishna Niyogi, and Matthew Traxler for use of their equipment. This work was supported by NIH grants R01GM114535 and R35GM139633 to M.E.T. We acknowledge that this work was conducted on the ancestral and unceded land of the Ohlone people.

## Author contributions

Conceptualization, K.J.K., F.J.W., K.C.M., and M.E.T.; Methodology, K.J.K., F.J.W., O.M.S., K.C.M., and M.E.T.; Validation: R.R.P.; Investigation, K.J.K. and L.V.I.; Writing – Original Draft, K.J.K. and M.E.T.; Writing – Review & Editing, K.J.K., O.M.S., K.C.M., and M.E.T.; Funding Acquisition, M.E.T.; Resources, O.M.S. and K.C.M.; Supervision, M.E.T.

## Declaration of interests

The authors declare no competing interests.

## Legends for supplemental items

**Figure S1.**
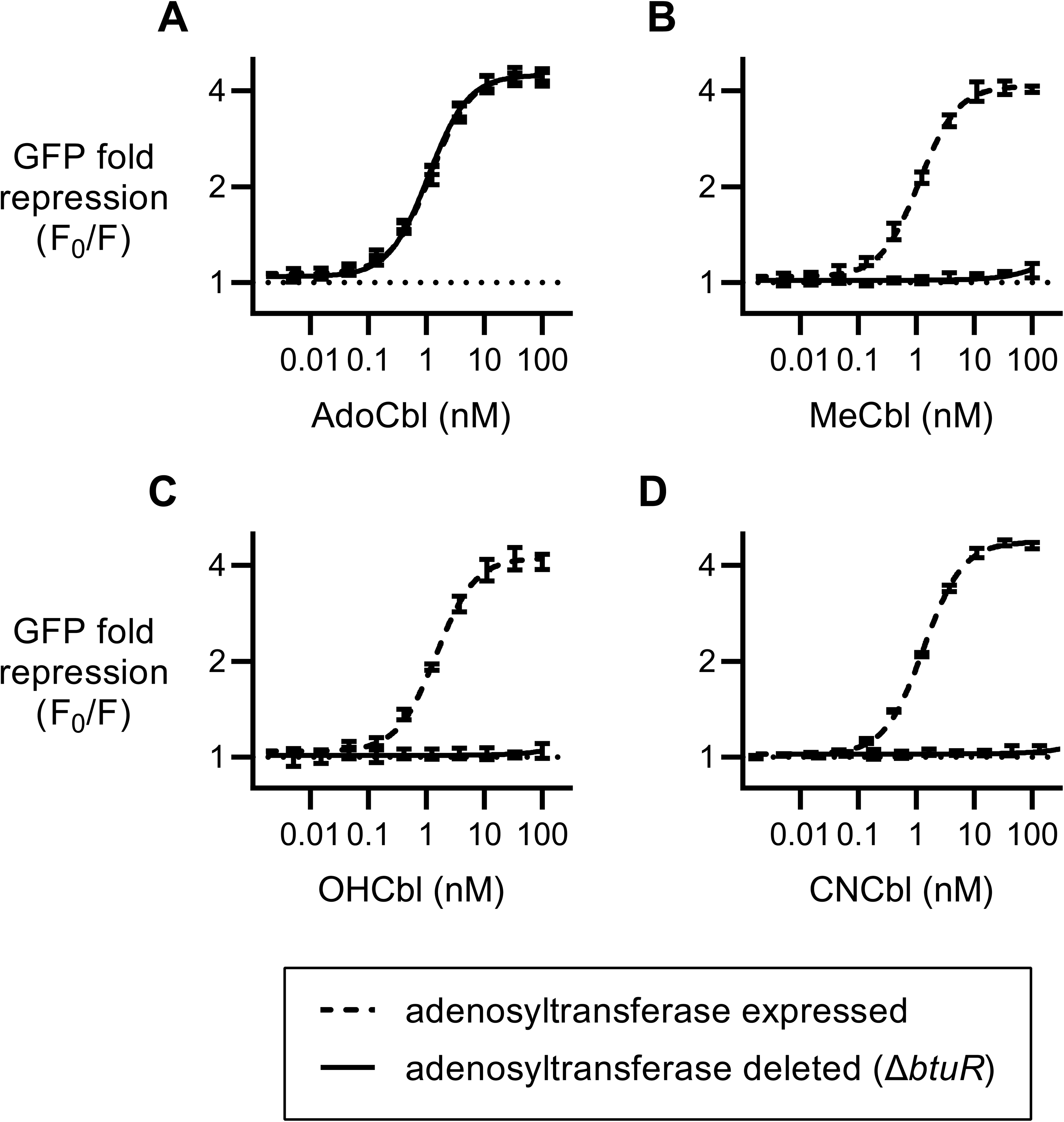
The *B. subtilis btuF* Cbl-riboswitch is sensitive to Cbl with 5’-deoxyadenosyl, but not methyl, hydroxyl, or cyanyl upper ligand moieties. Dose responses of *B. subtilis btuF* Cbl-riboswitch reporter strains with (A) adenosylcobalamin (AdoCbl), (B) methylcobalamin (MeCbl), (C) hydroxocobalamin (OHCbl), and (D) cyanocobalamin (CNCbl). Data points and error bars represent mean and standard deviation of four independent replicates. Horizontal dotted lines demarcate no change in expression. Note that the two dose-response curves in panel A are overlapping.

**Figure S2.**
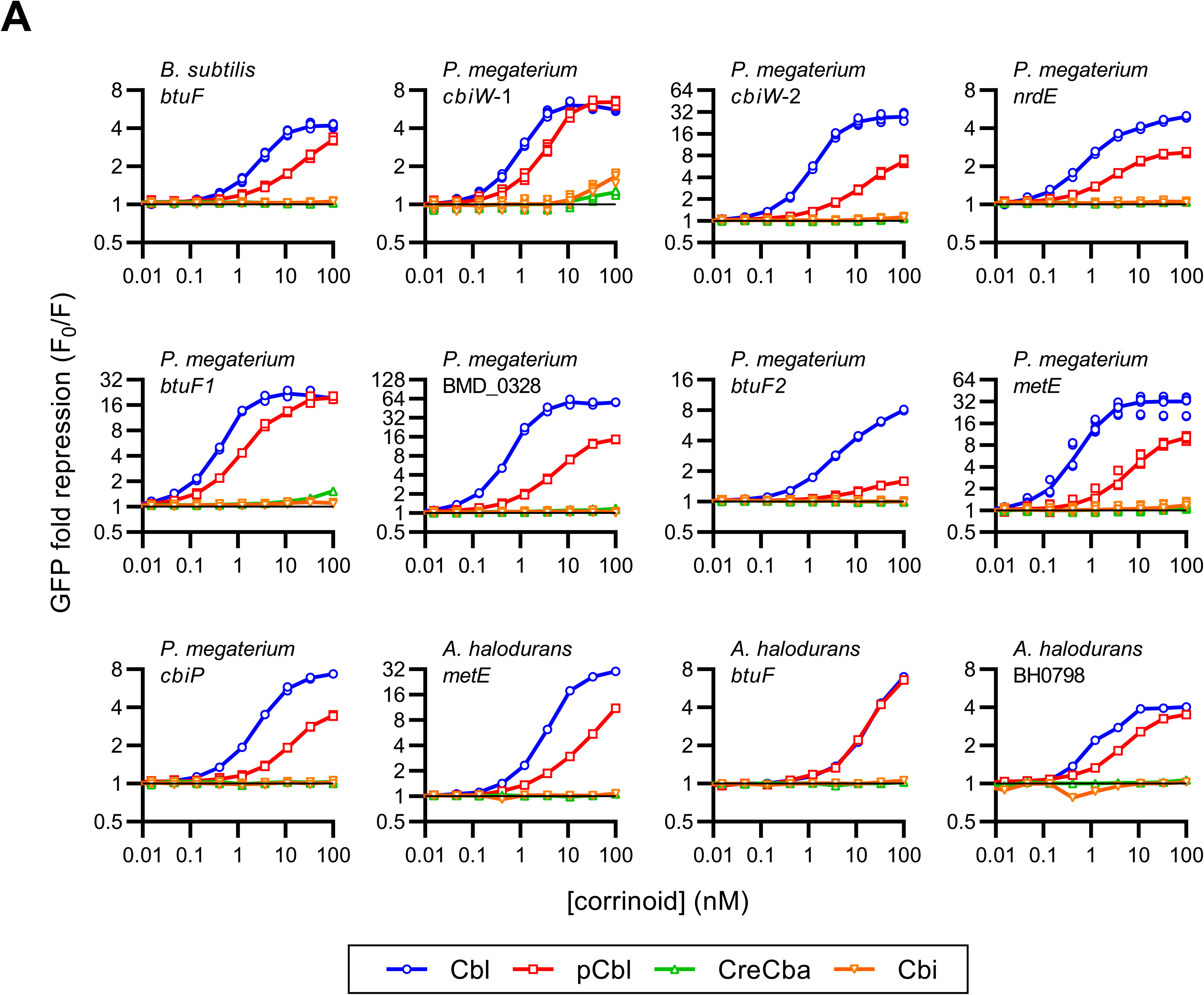
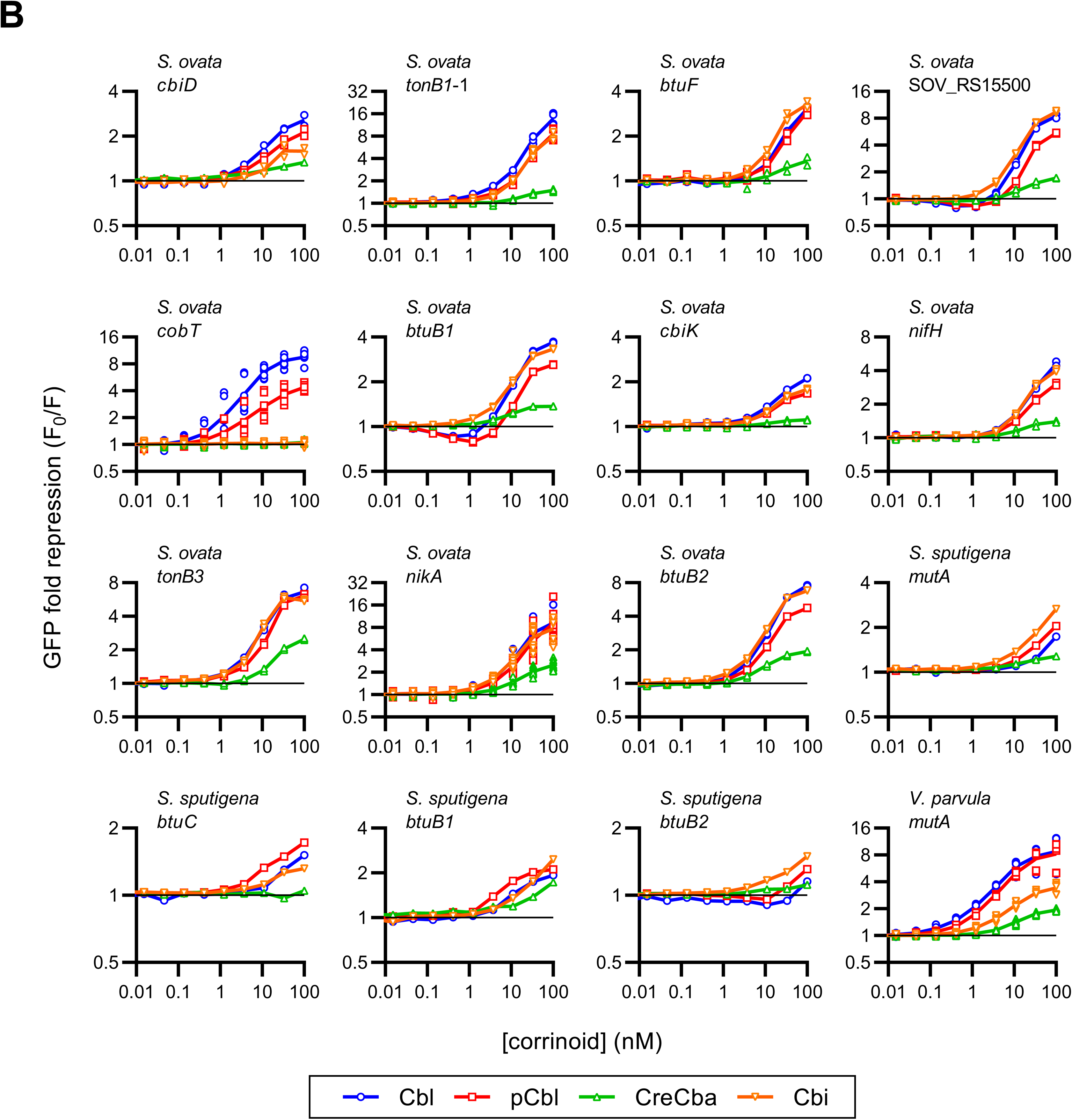
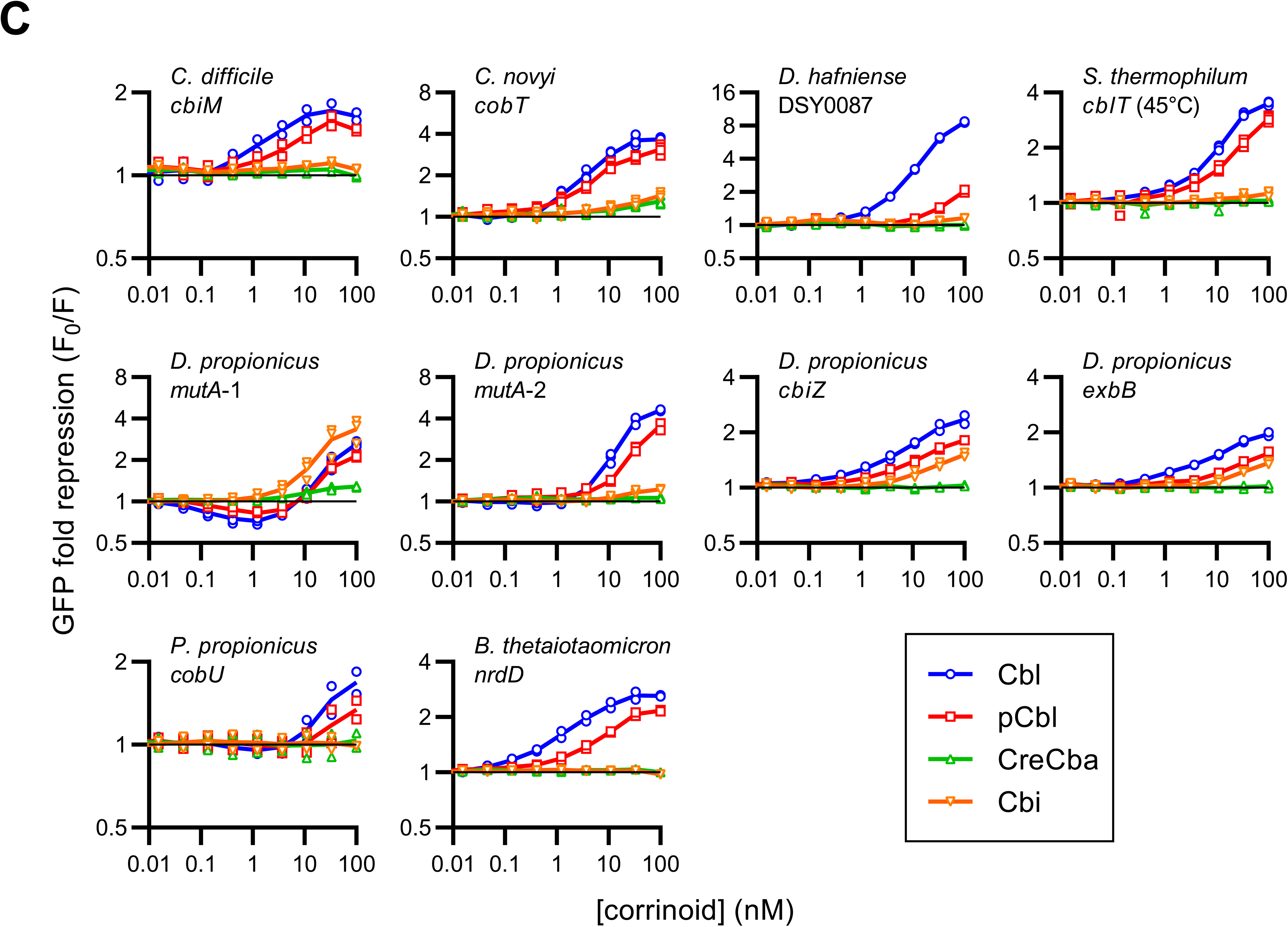
Corrinoid dose-responses of riboswitch reporter strains. Riboswitch names (species and gene downstream gene) are indicated for each dose-response. Data points are plotted for at least two independent experiments for each strain. Lines connect mean values. (A) Riboswitches from the class Bacilli. Species include *Bacillus subtilis*, *Priestia megaterium*, and *Alkalihalobacillus halodurans*. (B) Riboswitches from the class Negativicutes. Species include *Sporomusa ovata*, *Veillonella parvula*, and *Selenomonas sputigena*. (C) Riboswitches from the classes Clostridia, Deltaproteobacteria, and Bacteroidia. *Clostridioides difficile*, *Clostridium novyi*, *Desfulitobacterium hafniense*, and *Symbiobacterium thermophilum* are members of Clostridia. *Desulfobulbus propionicus* and *Pelobacter propoionicus* are members of Deltaproteobacteria*. Bacteroides thetaiotaomicron* is a member of Bacteroidia.

**Figure S3.**
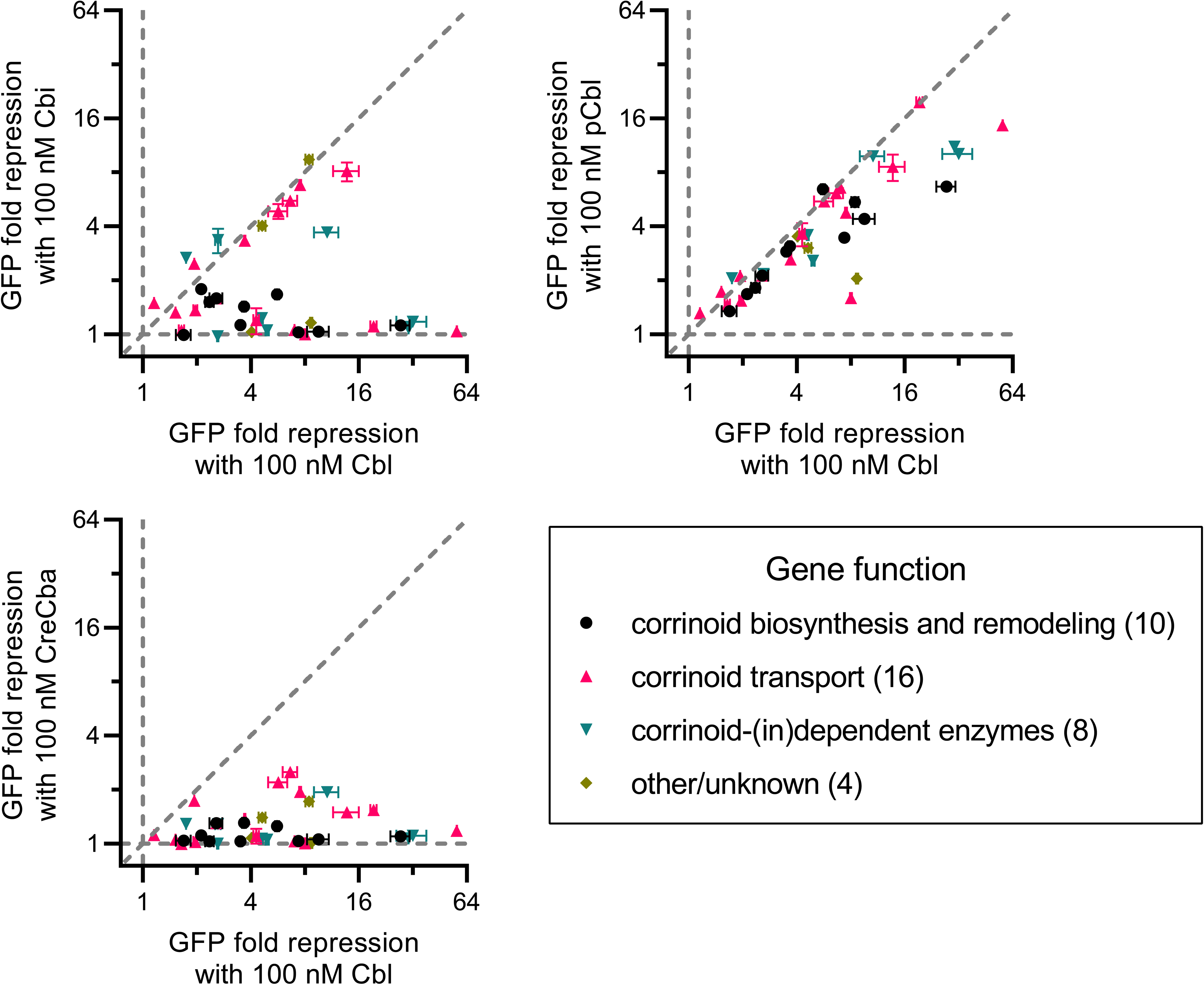
Cbl-riboswitch specificity does not cluster by regulatory gene target function. Pairwise comparisons of GFP fold repression induced by 100 nM doses of Cbl versus Cbi (top left) pCbl (top right), and CreCba (bottom left). Data points are colored by predicted function of downstream regulatory target genes. The number of riboswitches analyzed from each group is indicated in parentheses. Vertical and horizontal gray dashed lines demarcate a lack of response to one corrinoid. Diagonal line indicates equal repression between corrinoids. The distance of a point from the diagonal line indicates the bias in response towards one of the two corrinoids.

**Figure S4.**
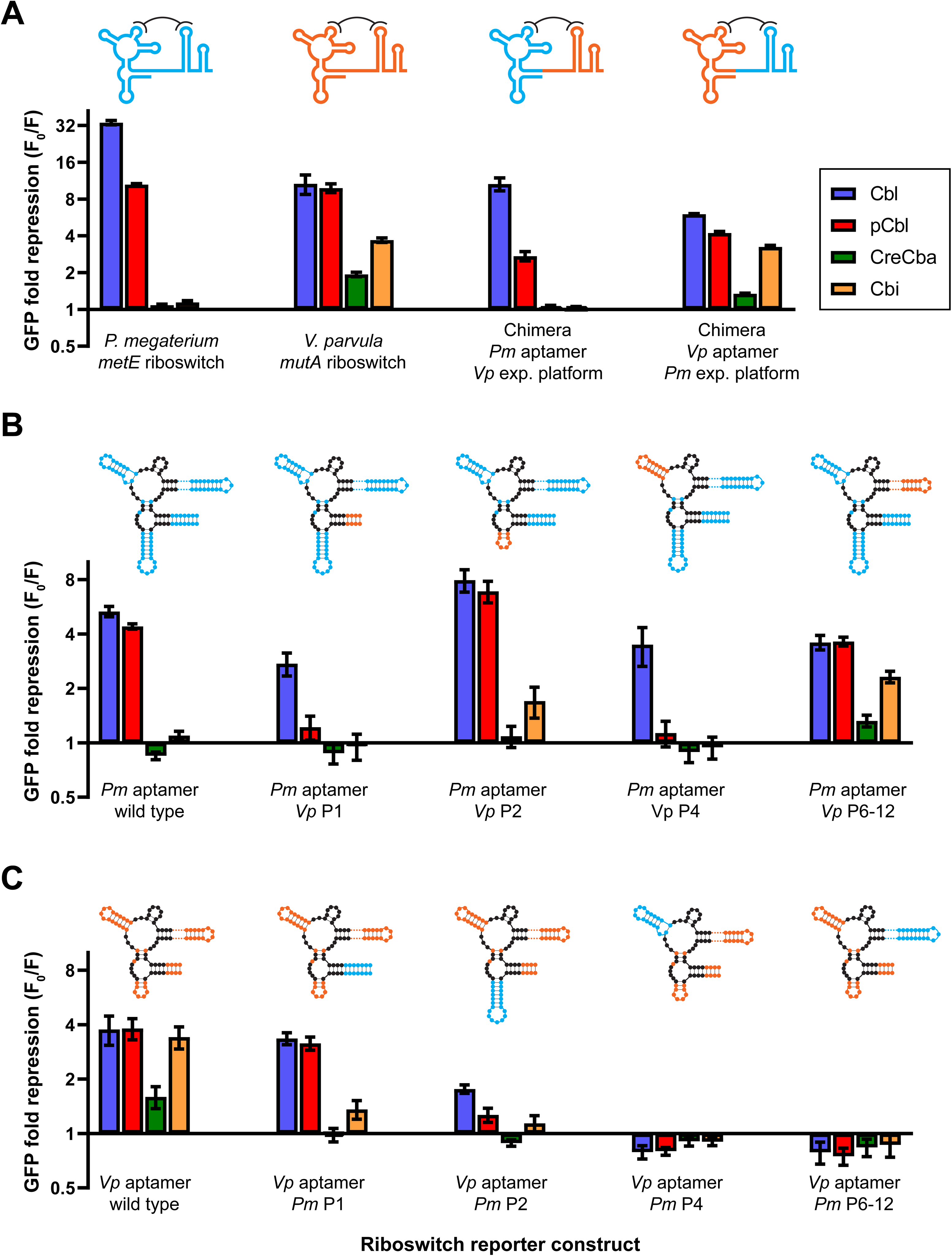
Chimeric riboswitches show that multiple components contribute to corrinoid specificity. Repression of GFP expression with 100 nM corrinoid is shown for (A) *P. megaterium metE* and *V. parvula mutA* riboswitches and chimeric fusions of their aptamer and expression platform domains. Aptamer subdomains swaps within the (B) *P. megaterium metE* riboswitch scaffold and (C) *V. parvul*a *mutA* riboswitch scaffold. Columns and error bars represent mean and standard deviation of 3 replicates. Cartoon riboswitches depict *P. megaterium* riboswitch sequence in cyan and *V. parvula* riboswitch sequence in orange. Sequences in black are identical between riboswitches.

**Figure S5.**
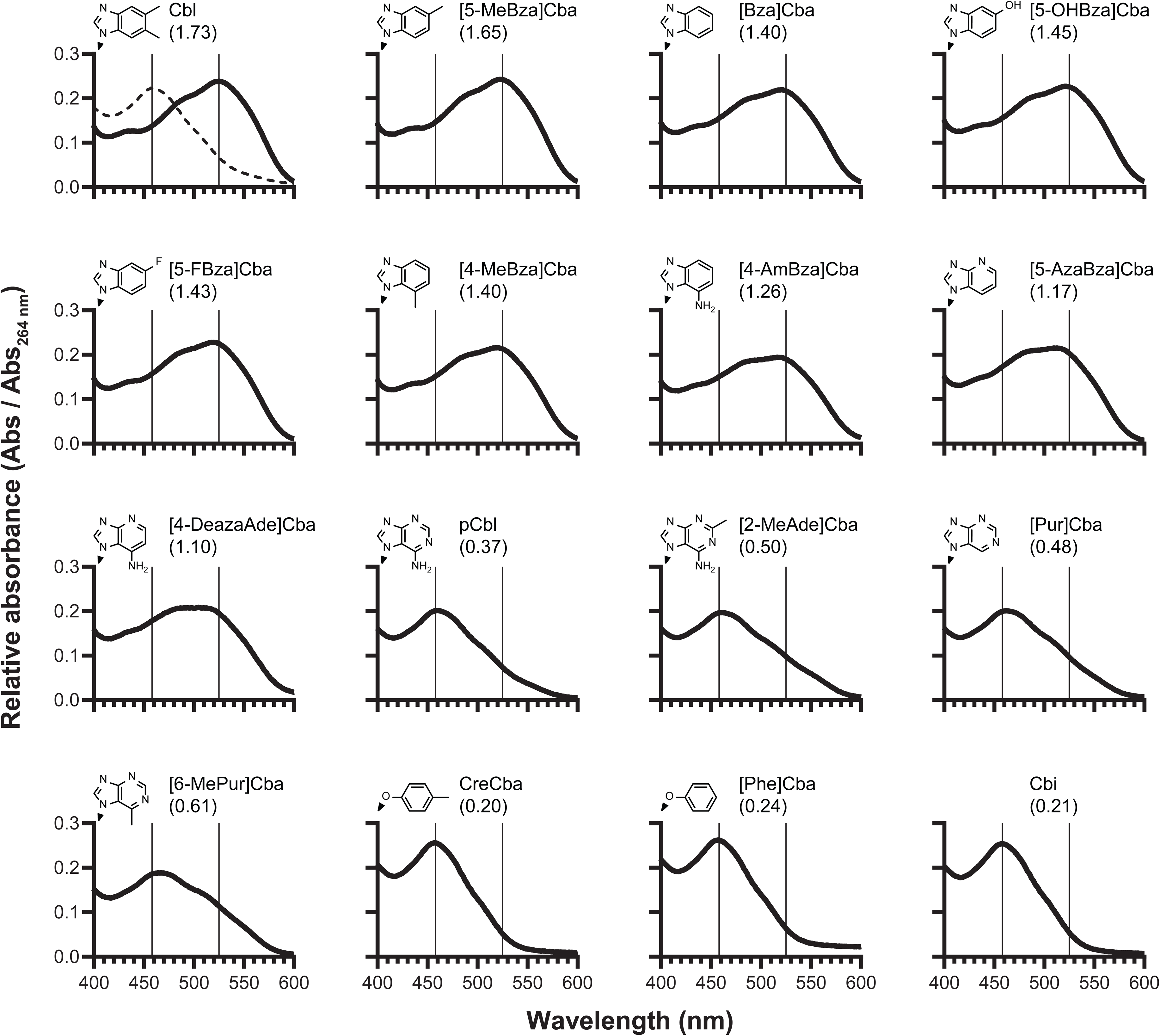
Corrinoid lower ligand structure impacts base-on tendency. Absorbance spectra of corrinoids were measured in a neutral buffered solution (pH 7.3). Absorbance peaks at 458 nm and 525 nm are associated with base-off and base-on conformations, respectively. Base-on tendency was measured as the ratio between absorbance at 525 nm and 458 nm (thin vertical lines). Each panel is labeled with corrinoid name, base-on tendency value (Abs_525nm_ / Abs_458nm_), and lower ligand chemical structure. Note that the tail structures of PheCba, CreCba, and Cbi cannot coordinate cobalt, and thus these corrinoids cannot be assigned a ‘base-on tendency’ per se. The Abs_525nm_/Abs_468nm_ ratios of PheCba, CreCba, and Cbi simply reflect an absence of any Co-N coordination at the α axial face. For reference, the dashed line in the upper left panel is the absorbance spectrum of nearly complete base-off Cbl measured in acidic solution (pH 1.57).

**Figure S6.**
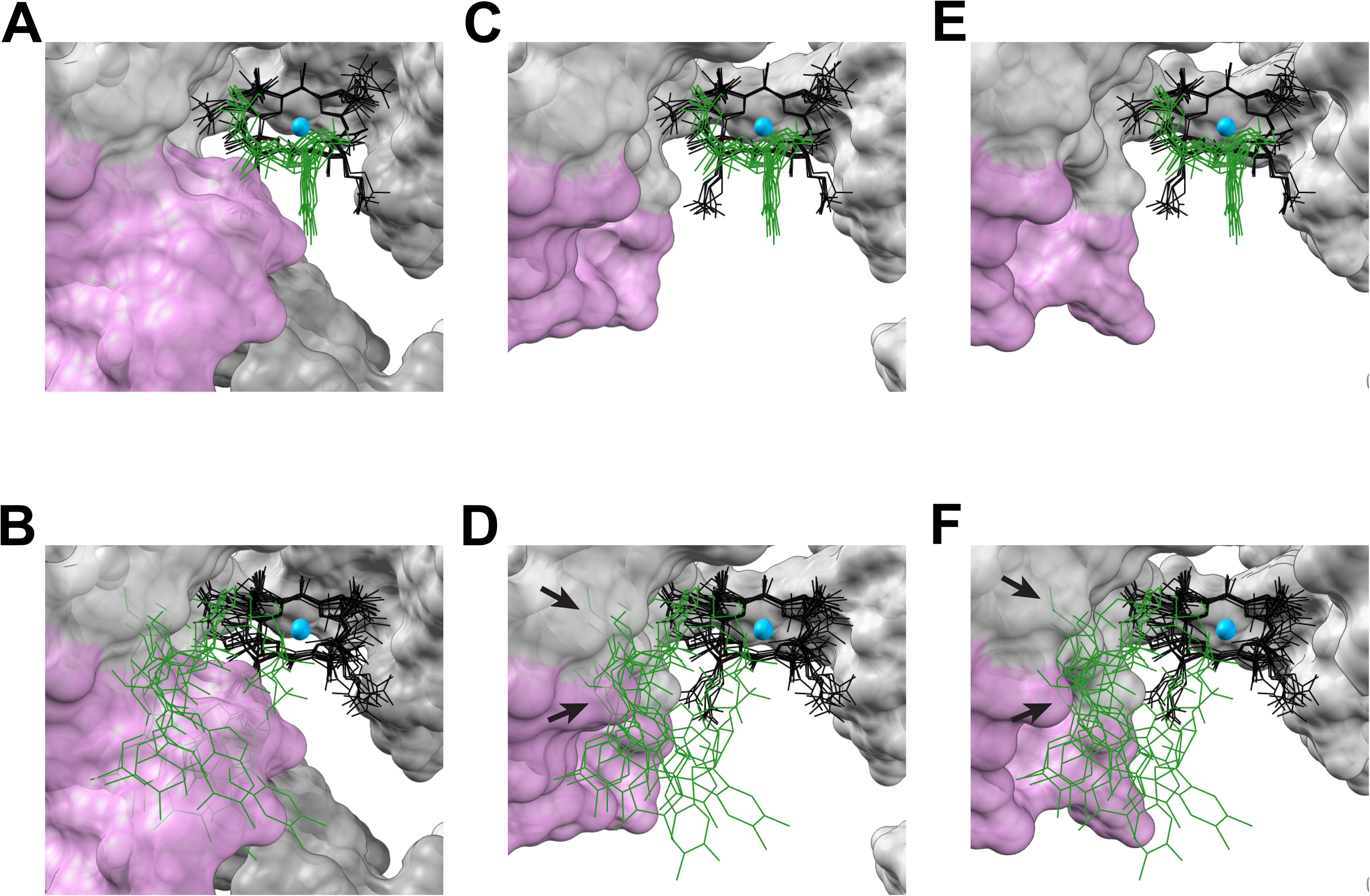
Cbl-binding sites in X-ray crystal structures of various Cbl-riboswitches. The marine metagenome derived riboswitch *env8* (PDB ID 4FRN) (A, B), the *B. subtilis btuF* Cbl-riboswitch (PDB ID 6VMY) (C, D), and the *S. thermophilum cblT* Cbl-riboswitch (PDB ID 4GXY) are depicted with base-on (A, C, E) and base-off (B, D, F) Cbl alignments. Cbl models were aligned by the cobalt and coordinating nitrogen atoms in the corrin ring. Structures of the corrin ring, cobalt, and tail of Cbl are colored in black, blue and green, respectively. Upper ligand structures of Cbl were omitted for clarity. Riboswitch RNA structures are depicted as space-filled models with the L5-L13 kissing loop in pink in panels A and B, and just the L5 in pink in panels C-F. The rest of the RNA structures in each panel are in gray. Arrows in panels D and F point to clashes between RNA and the Cbl tails.

**Figure S7.**
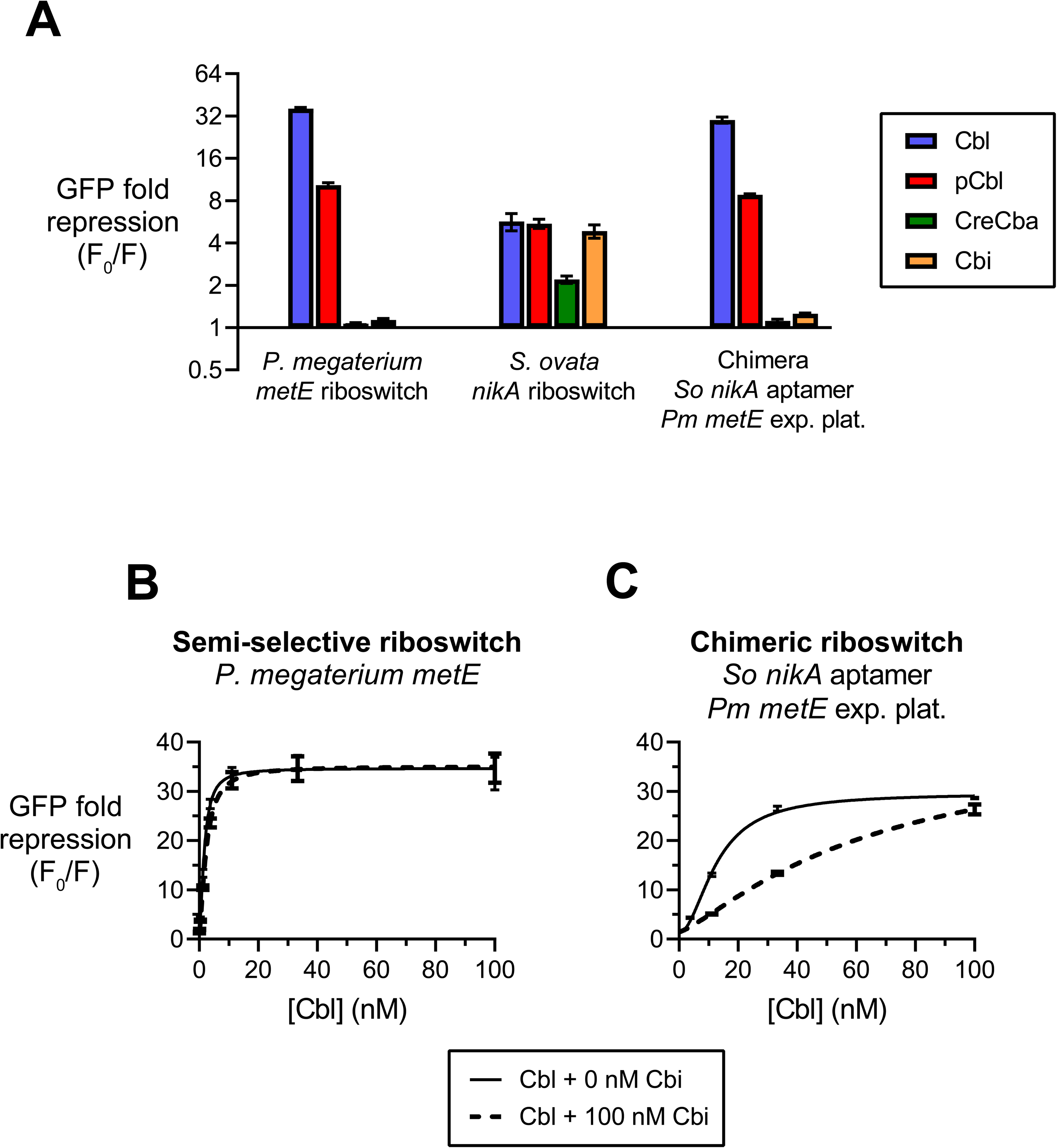
The aptamer and expression platform of the *P. megaterium* riboswitch contribute to corrinoid selectivity. (A) Repression of GFP expression with 100 nM corrinoid is shown for the *P. megaterium metE* riboswitch, *S. ovata nikA* riboswitch, and a chimeric fusion of the *S. ovata nikA* aptamer and *P. megaterium metE* expression platform domains. Cbl dose-responses with or without 100 nM Cbi in (B) *P. megaterium metE* and (C) chimeric riboswitches. Data points and error bars represent mean and standard deviation of four independent replicates.

**Table S1 – Strain list and riboswitch information.** (A) List of strains used in this study. (B) Genome and sequence information of wild type riboswitches analyzed in this study. (C) Sequences of mutant riboswitches analyzed in this study.

**Table S2 – X-ray crystal structural models used for 3D Cbl structural alignments.**

**Data S1 – Riboswitch sequence alignment in Stockholm sequence format.**

## Notes

### Competing Interest Statement

The authors have declared no competing interest.

### Summary of Updates

Revised text for readability. Added new topics to the Discussion section. Merged Figure S10 with Figure 6. Merged Figures S2, S3, and S4. Merged Supplemental file 1 and Supplemental file 2. Changed numbering of supplemental items. Added a few relevant citations.

