## Supplementary material for "Cobalamin riboswitches are broadly sensitive to corrinoid cofactors to enable an efficient gene regulatory strategy": Table S2

| PDB ID | Bound macromolecule | Organism | Base-on/off |
| --- | --- | --- | --- |
| 4GMA | Cbl-riboswitch | *Thermanaerobacter tengcongensis* | Base-on |
| 4FRN | Cbl-riboswitch | Unknown (marine metagenome sequence) | Base-on |
| 4FRG | Cbl-riboswitch | Unknown (marine metagenome sequence) | Base-on |
| 4GXY | Cbl-riboswitch | *Symbiobacterium thermophilum* | Base-on |
| 6VMY | Cbl-riboswitch | *Bacillus subtilis subsp. subtilis* str. 168 | Base-on |
| 1ET4 | Synthetic RNA aptamer | Not applicable | Base-on |
| 4REQ | Methylmalonyl-CoA mutase | *Propionibacterium freudenreichii subsp. shermanii* | Base-off |
| 2XIJ | Methylmalonyl-CoA mutase | *Homo sapiens* | Base-off |
| 1K7Y | Methionine synthase | *Escherichia coli* | Base-off |
| 5D08 | Epoxyqueuosine reductase | *Bacillus subtilis subsp. subtilis* str. 168 | Base-off |
| 5D6S | Epoxyqueuosine reductase | *Streptococcus thermophilus* | Base-off |
| 5C8D | CarH photoreceptor | *Thermus thermophilus* HB27 | Base-off |
| 4RAS | Reductive dehalogenase | *Nitratireductor pacificus* pht-3B | Base-off |
| 5CJU | Isobutyrl-CoA mutase | *Cupriavidus metallidurans* CH34 | Base-off |
| 1I9C | Glutamate mutase | *Clostridium cochlearium* | Base-off |
| 1XRS | D-lysine 5,6-aminomutase | *Acetoanaerobium sticklandii* | Base-off |

**Table S2 – X-ray crystal structural models used for 3D Cbl structural alignments.**
